# Detecting nanoscale distribution of protein pairs by proximity dependent super-resolution microscopy

**DOI:** 10.1101/591081

**Authors:** Alexander H. Clowsley, William T. Kaufhold, Tobias Lutz, Anna Meletiou, Lorenzo Di Michele, Christian Soeller

## Abstract

Interactions between biomolecules such as proteins underlie most cellular processes. It is crucial to visualize these molecular-interaction complexes directly within the cell, to show precisely where these interactions occur and thus improve our understanding of cellular regulation. Currently available proximity-sensitive assays for in-situ imaging of such interactions produce diffraction-limited signals and therefore preclude information on the nanometer-scale distribution of interaction complexes. By contrast, optical super-resolution imaging provides information about molecular distributions with nanometer resolution which has greatly advanced our understanding of cell biology. However, current co-localization analysis of super-resolution fluorescence imaging is prone to false positive signals as the detection of protein proximity is directly dependent on the local optical resolution. Here we present Proximity-Dependent PAINT (PD-PAINT), a method for sub-diffraction imaging of protein pairs, in which proximity detection is decoupled from optical resolution. Proximity is detected via the highly distance-dependent interaction of two DNA labels anchored to the target species. Labeled protein pairs are then imaged with high contrast and nanoscale resolution using the super-resolution approach of DNA-PAINT. The mechanisms underlying the new technique are analyzed by means of coarse-grained molecular simulations and experimentally demonstrated by imaging DNA-origami tiles and epitopes of cardiac proteins in isolated cardiomyocytes. We show that PD-PAINT can be straightforwardly integrated in a multiplexed super-resolution imaging protocol and benefits from advantages of DNA-based super-resolution localization microscopy, such as high specificity, high resolution and the ability to image quantitatively.

## Introduction

Characterizing protein interactions by detection of protein-protein complexes is the basis of understanding many processes in biology.^1^ Often, these are detected by in-vitro methods such as co-immunoprecipitation, cross-linking or affinity blotting.^2,3^ It is increasingly evident that besides detecting the mere presence of protein-protein interactions, it is important to determine where these occur within a cell or tissue, since the nanoscale organization of signaling complexes directly controls cell function.^4,5^ To this end, methods have been developed that are based on labeling the features of interest with synthetic DNA oligonucleotides, conjugated to antibodies or other molecular markers. The oligonucleotides act as proximity probes, and a subsequent amplification step is implemented to produce a fluorescent signal detectable by a conventional microscope. In an (in-situ*) proximity ligation assay* (PLA), enzymatic amplification occurs via the rolling circle method,^5–8^ while in the ProxHCR scheme amplification is non-enzymatic and relies on a hybridization chain reaction.^9,10^ However, the high fluorescent amplification in both methods also effectively restricts them to diffraction-limited imaging. This implies that the presence of protein pairs can be located to within a certain region but generally precludes accurate quantification and visualization of their distribution at the nanometer scale.^11^

An alternative approach to detecting protein-protein interactions relies on a FRET-based assay, a fluorescent method that is distance sensitive in the nanometer range.^12^ While FRET can sensitively detect the presence of protein-protein interactions, FRET imaging implementations are generally diffraction-limited and therefore it is not possible to resolve the distribution of protein pairs with true molecular resolution.

Imaging with spatial resolution beyond the diffraction limit can now be achieved by a wide range of super-resolution techniques, such as structured illumination microscopy (SIM),^13^ stimulated emission depletion (STED) microscopy,^14,15^ (fluorescence) photoactivated localization microscopy ((F)PALM)^16,17^ or (direct) stochastic optical reconstruction microscopy ((d)STORM).^18,19^ DNA-PAINT (Point Accumulation Imaging in Nanoscale Topography) is a super-resolution imaging technique that relies on oligonucleotide interactions, a property that it shares with PLA and ProxHCR.^20^ In DNA-PAINT, the proteins of interest are labeled with a short DNA oligonucleotide, or “docking” strand. The transient binding and consequent brief immobilization of fluorescently labeled complementary “imager” strands enables detection as a fluorescent “blink” and precise localization. The accumulated localization data is used to reconstruct a super-resolution image.

Fluorescent super-resolution techniques can be used to acquire multi-target images which, in principle, allow estimating the proximity of protein targets by fluorescence co-localization. However, unequivocal identification of protein-protein pairs is complicated by the fact that co-localization can be prone to false-positive signals, as the precision of co-localization is directly dependent on the local imaging resolution. Resolution can vary considerably, especially in optically complex samples such as thick cells or tissue sections. Often, there is limited resolution along the optical axis (several hundreds of nanometers in 2D super-resolution techniques, >40 nm in 3D methods)^21^ which can lead to additional false positives. Furthermore, registration errors between multiple channels, e.g. due to chromatic aberrations or sample drift, can lead to incorrect co-localization estimates.

Here, we introduce a proximity-sensitive super-resolution imaging method which allows detection of the presence and characterization of the local density and nanoscale distribution of protein-protein complexes, determining the detailed structure and morphology of possible clusters. Importantly, the proximity detection is fully decoupled from local imaging resolution and optical multi-channel registration is not required. In our approach, two proteins of interest are labeled with DNA constructs, which are designed to interact when in molecular proximity, and as a result allow for a DNA-PAINT signal to be detected. One construct includes a DNA-PAINT docking sequence, which however is protected by a stable DNA stem loop structure, rendering the docking site effectively inaccessible. If the second target is within a distance of approximately 10 to 20 nm (depending on the specific design, see below), the second DNA construct associated with it can displace the stem of the hairpin, thereby unfolding the loop, fully exposing the docking site, and enabling DNA-PAINT imaging of the structure. We term this new approach *Proximity-Dependent PAINT* (PD-PAINT) and prove its functionality by means of coarse-grained molecular simulations and experimentally by imaging DNA-origami tiles with pre-designated spacings. We then demonstrate the applicability of PD-PAINT to an important biological scenario, namely the imaging of epitopes on cardiac ion channels and nearby supporting proteins.

PD-PAINT combines proximity detection with all the advantages of DNA-PAINT. These include high specificity, straightforward implementation on a conventional fluorescence microscope, freedom in the choice of dye and wavelength and importantly the possibility of readily achieving high photon yields, resulting in very high resolution.^20,22^ In addition, by exploiting the principle of quantitative PAINT (qPAINT), PD-PAINT enables accurate determination of the local density of protein pairs in the sample.^20,23^ Finally, PD-PAINT is fully compatible with multi-color imaging and can be implemented in addition to conventional DNA-PAINT of multiple other channels, essentially without performance penalty. This is demonstrated here by following an Exchange-PAINT protocol,^24^ to image the protein complex as well as the individual protein epitopes in separate channels.

## Results and Discussion

Fig. 1 shows the principle of PD-PAINT, where fluorescence signals are only detected if two epitopes of interest are close to each other. To implement PD-PAINT, the two target epitopes are labeled with nanostructures S1 and S2 via suitable markers. Labeling can be done by linking the nanostructures to antibodies (or nanobodies), or by hybridizing them to other DNA strands, previously connected to the epitopes. Strand S1 contains the 9 nt docking sequence consisting of domains 5* (3 nt) and 6 (6 nt). The docking sequence is complementary to the imager strand P1 (6*-5). However, when S1 is spatially isolated, the docking site is protected by a closed stem-loop motif in which 5* is hybridized to the complementary domain 5, and 6 is bent to form a short loop. The loop is further stabilized by the complementary domains 3-4 and 3*-4* (11 bp). In the closed-loop configuration of S1 the docking site has negligible affinity for the P1 imager (Fig. 1a), as previously determined in the context of catalytic DNA reactions^9^ and further demonstrated here by means of coarse-grained computer simulations discussed below. The second target is labeled with nanostructure S2, complementary to the stem region of S1 (3-4) as well as the un-protected toe-holding domain 1-2. The latter would promote rapid S1-S2 dimerization, should the two nanostructures be present in sufficient concentration and have unrestricted access to each other, e.g. during labeling steps. To prevent unwanted S1-S2 dimerization, S2 is therefore initially protected by a shield strand B, which occupies the toe-holding region 1*-2* and is further stabilized by adjacent complementary domains 3 and 7. Once both S1 and S2 are bound to their appropriate targets, and any excess washed away, a removal strand R is used to displace B from S2, freeing 1*-2* (Fig. 1b). At this stage, S1-and S2 can dimerize, but only if their tethering locations are within a given maximal distance from each other. When S1 and S2 are fully hybridized, the open loop exposes the docking sequence 5*-6 of S1, allowing transient binding of the complementary imager P1, which results in frequent detection of these binding events as fluorescent blinks, i.e., a DNA-PAINT signal (Fig. 1c). Whereas Fig. 1a-c show a direct tethering geometry (geometry 1), in the experiments described below an indirect tethering geometry (geometry 2) was employed, relying on two attachment strands D1 and D2 which are anchored on the substrate. S1 and S2-B, when added in solution, attach to D1 and D2, respectively, as shown in Fig. 1d.

**Figure 1.**
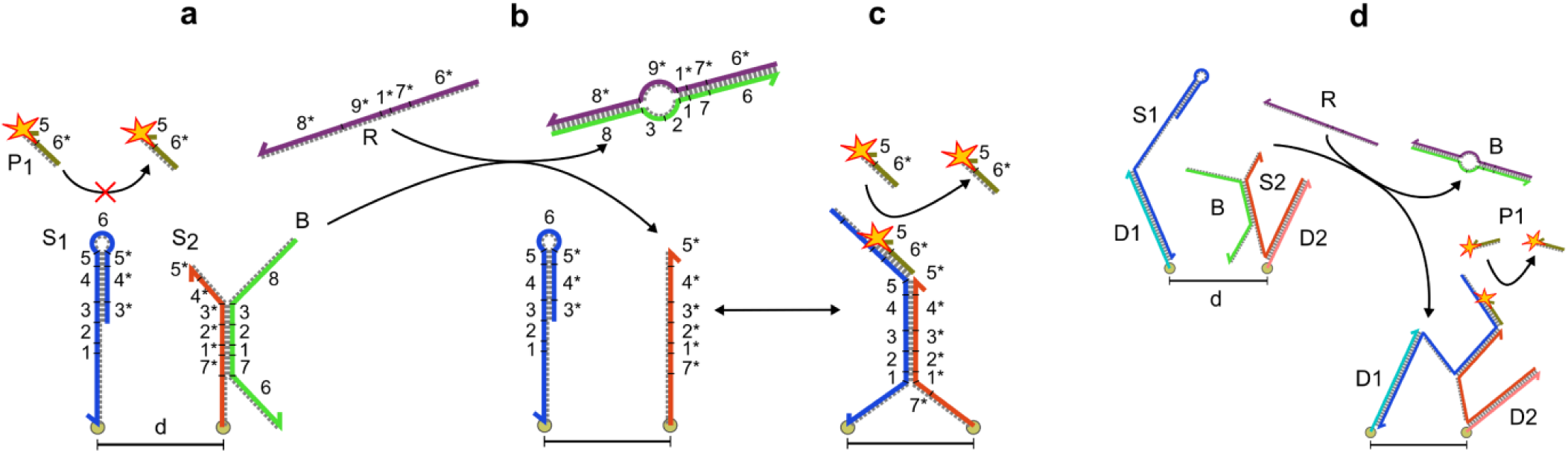
Principle of Proximity-Dependent PAINT (PD-PAINT). **a:** Strand S1 forms a stem loop structure which prevents imagers (P1) binding to the docking domain (5*-6), due to a high curvature of domain S1-6 and hybridization of domain S1-3*-5* to S1-3-5. Strand S2 contains domains 1*-5* which are complementary to 3-5 of the S1 stem and the unbound toehold region 1-2 on S1. The corresponding toe-hold domain 1*-2* on S2, along with neighboring domains 7* and 3*, are initially protected by shield strand B in order to prevent unwanted S1-S2 dimerization during the sample functionalization stages. **b:** Addition of the shield remover R strips the shield B, and, should S1 and S2 be in close proximity, triggers their dimerization initiated by the 1-2 toe-hold, which results in the opening of the S1 loop. **c:** The now exposed docking domain 5*-6 allows the transient binding of an imager to the S1-S2 complex so that a super-resolution image can be obtained by DNA-PAINT. **d:** Alternative tethering geometry (geometry 2) in which strands S1 and S2 are attached to the substrate via attachment strands D1 and D2. In this configuration the double stranded domain formed by D1 and S1 can rigidly rotate around the attachment point of D1 whereas the flexible single-stranded section of S2 is free to move around the attachment point of D2.

To explore the functionality of the proposed PD-PAINT scheme we investigated its thermodynamic properties by means of Monte Carlo (MC) simulations using the oxDNA coarse-grained model of nucleic acids, as described in detail in the Supporting Information.^25,26^ For PD-PAINT to elucidate the co-localization of epitopes, it must obey two thermodynamic design criteria. First, the S1 loop must open if S2 is nearby. Second, the closed loop should not interact with its imager in a way that leads to detectable blinks, i.e. any transient binding event of P1 to the closed S1 loop must be much briefer than those detected for the open S1 loop.

We implemented free-energy calculations to determine the likelihood of the formation of S1-S2 dimers, resulting in the opening of the S1 loop, as a function of the distance *d* between their anchoring points. Simulations were performed for both direct anchoring (geometry 1, Fig. 1 a-c) and indirect anchoring (geometry 2, Fig. 1d). In geometry 1, the reactive domains of S1 and S2 are separated from their anchoring points by a 15-nt polyThymine sequence, while in geometry 2, directly relevant to experiments, the reactive domains of S1 are further spaced by a 32 base-pair rigid double-stranded (ds) DNA domain.

Figure 2a reports the computed S1-S2 dimerization free energy as a function of the tethering distance. In both tested geometries the reduction in free energy of dimerization is substantial at short separations, with minima between -12 and -10 kBT. At large separation distances the dimerization free energy sharply rises, with the increase occurring at larger separations for geometry 2 (red symbols), due to the additional 32bp dsDNA spacer connected to S1. The S1-S2 dimerization probability is shown in Fig. 2b. Both geometries display an approximately step-like response, with dimerization probability essentially equal to one for separations below ∼10 nm for geometry 1 (blue symbols) and ∼20 nm for geometry 2 (red symbols). At larger separation distances, dimerization rapidly becomes essentially impossible, and the probability falls to zero at ∼16 nm (geometry 1) and ∼25 nm (geometry 2).

**Figure 2.**
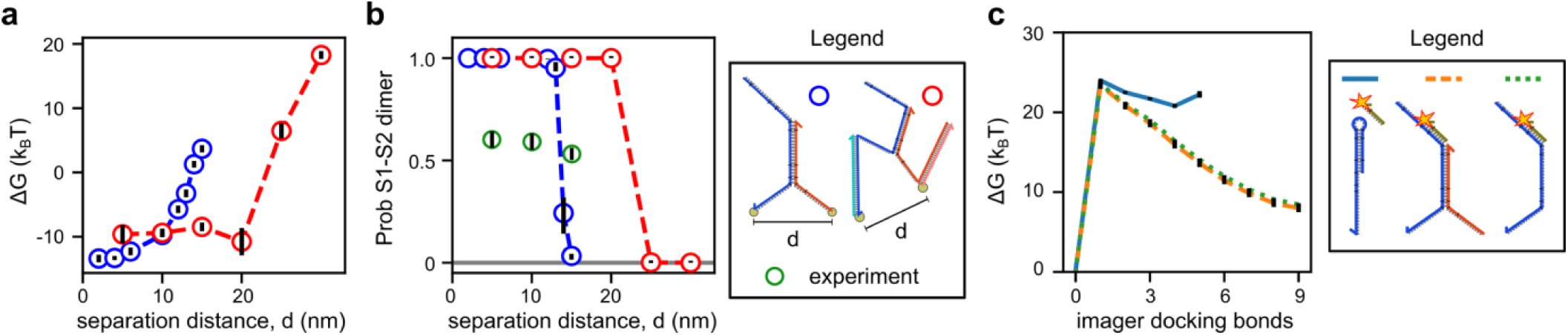
Thermodynamic properties of PD-PAINT from coarse-grained simulation. **a:** The free energy change of S1-S2 hybridization is sensitive to the separation distance between tethers and to the structure connecting the tether to the loop or its complement. At close inter-tether distances (< 5 nm), hybridization is strongly favored for both tested tethering geometries. At higher distances, hybridization free energy increases monotonically for the direct tethering geometry (geometry 1, blue) due to stretching of the entropic spring formed by the two 15 polyThymine linkers. For the indirect-tethering geometry (geometry 2, red), the 32 base pair dsDNA spacer reduces the dependency of free energy on inter-tether distance. **b:** Simulated dimerization probabilities are close to 1 at inter-tether distances shorter than 10 nm (geometry 1), or 20 nm (geometry 2). At greater distances (∼16 nm for geometry 1 and ∼25 nm for geometry 2) dimerization probability sharply transitions to ∼0. Green circles indicate experimental dimerization probabilities as extracted from the kinetic analysis of blinking events recorded on origami test samples, specifically from a numerical fit of the dark-time distributions to the outcomes of explicit Markov simulations (Fig. 4, SI text, Fig. S2, and Table S5). Note that simulations are found to overestimate dimerization probability. **c:** Hybridization of the imager to S1 as a closed loop is inhibited in the absence of S2 and restored by the presence of S2. Free energy profiles for formation of Watson-Crick bonds between the imager and S1 are shown, relative to the unhybridized state. We compare the closed loop state of S1 (blue line), the open state after hybridization of S2 (orange dashed) and conventional DNA-PAINT binding (green dotted). Note that closed loop states with >5 bonds resulted in large free energies whose contribution to the overall hybridization free energy is negligible. Legend: the three configurations tested in panel **c**. All DNA sequences used are shown in Table S6. All error bars are given as one standard error.

For the presence and location of S1-S2 dimers to be positively identifiable via DNA-PAINT, binding of the imager to closed S1 loops must be practically undetectable, yet the imager must have a sufficient affinity for the exposed docking site when the S1 loop is open. We verify these requirements by estimating the interaction free-energy between the P1 imager and S1 in its closed and open loop states, shown in Fig. 2c as a function of the number of formed base-pairing bonds between the two strands. The free energy barrier for the formation of the first bond is similar between the open and closed loop configurations but while in the closed-loop case further base pairing does not result in a significant drop in free energy, a steep monotonic decrease is observed for the open-loop configuration. As a result, we estimate the average duration of binding events between the imager to the closed S1 loop as ∼2 µs, well below the detection threshold for a typical DNA PAINT experiment (see Supporting Information and Table S9). In turn, for the open loop configuration we predict the binding events to last ∼0.2 – 0.3 s, ideal for DNA-PAINT at typical frame integration times of 50-300 ms (Table S9). Comparison between the binding free energy of P1 to the open S1 loop of a S1-S2 dimer to that towards a conventional single-stranded (ss) DNA docking strand reveals no perceptible difference (Fig. 2c), and, consistent with this observation, the predicted bond lifetime in the two cases is comparable (Table S9).

Having established the viability of our platform in-silico, we experimentally characterized the PD-PAINT scheme on DNA-origami test samples. DNA origami tiles were produced following a standard protocol,^20^ as described in detail in the Supporting Information. The tiles featured 6 pairs of single stranded D1 and D2 overhangs, to which S1 and S2 can hybridize as illustrated in Fig. 3, thus realizing the conditions of geometry 2 in Fig. 1c. Three types of tiles were tested, with nominal distances between S1 and S2 tethering points *d*=5, 10 or 15 nm (Fig 3c-e). Strand D1 contains a 5*6 docking domain, complementary to imager P1 (Fig. 3a), which allowed us to perform conventional DNA-PAINT imaging in the absence of S1 and S2, and then compare it with PD-PAINT under identical experimental conditions (in separate experiments we also determined the addressability of D2, see Fig. S1). All PD-PAINT experiments started with this direct imaging of D1, an experimental stage we refer to as the “direct phase” (Fig. 3a-i).

**Figure 3.**
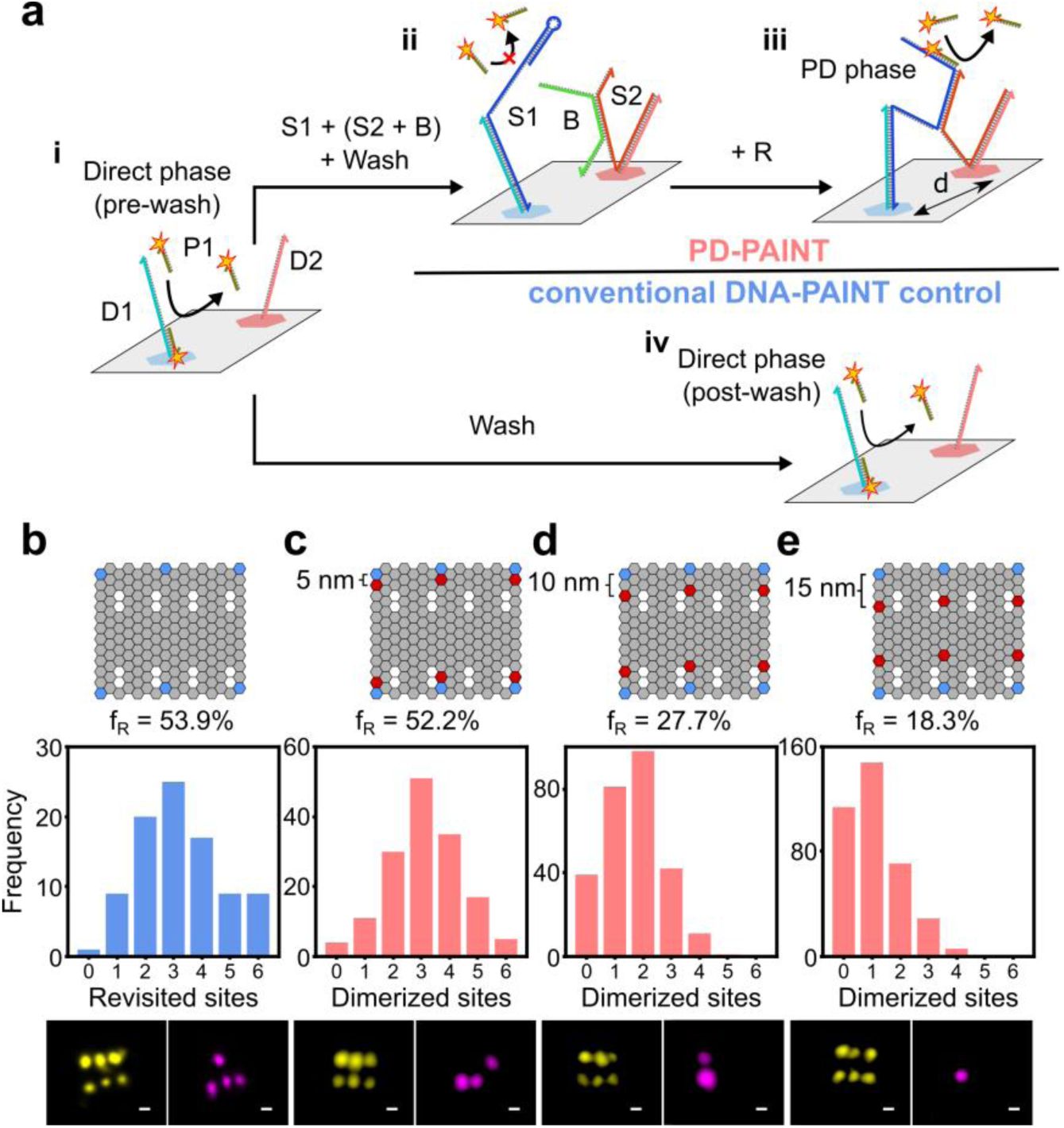
Experimental assessment of PD-PAINT with DNA origami test samples. **a:** Origami tiles with fixed D1-D2 separation *d*=5, 10 or 15 nm were initially probed with imager P1 against a complementary domain in D1 (direct phase, i). Strand S1 was then added, which binding to D1 prevents its direct imaging, followed by the S2-B construct binding to D2 (ii). Introduction of the shield removal sequence R displaces B from S2, enabling S1-S2 dimerization, the exposure of the P1 binding site on S1, and the detection of PD-PAINT signals (PD phase, iii). In control samples, buffer exchange (washing) steps were performed without the addition of PD-PAINT probes, before a second (post-wash) direct imaging phase (iv). **b-e:** assessment of the percentage of detectable origami sites retained between the (pre-wash) direct phase and either the post-wash direct phase (control experiment, **b**) or the PD phase (PD-PAINT experiments with increasing *d*, **c-e**). The fraction of retained sites fR is extracted by considering all origami tiles displaying six visible sites in the (pre-wash) direct phase. The distribution of the number of sites per tile retained in the post-wash direct phase or the PD-phase is shown in the histograms. Example renderings of origami imaged in the pre-wash direct phase (left) and then in the post-wash direct phase / PD phase (right) are shown below. The number of tiles sampled was 90, 153, 272 and 369 for panels **b**-**e**, respectively. Scale bars: 30 nm.

After completing the direct phase, S1 was added, which binds to the entire D1 strand thus blocking access to the previously imaged docking domain on D1 (Fig. 3a, top). Subsequently we included the protected S2 nanostructure (S2-B), which binds to overhang D2. Finally, the shield strand B was removed by adding an excess of remover strand R, exposing the S1-S2 toehold domain and enabling their dimerization. The latter renders the docking domain on S1 accessible for imaging, an experimental stage that we indicate as “PD phase”. A complete time trace of the experimental protocol and the resulting event rates in a typical experiment is shown in Fig. S5.

For further assessments we also performed control experiments in which the (pre-wash) direct phase is followed by buffer exchange steps similar to those required to introduce the PD-PAINT constructs, but without actually adding S1 and S2. Instead, direct imaging of D1 is resumed with conventional DNA-PAINT in a “post-wash direct phase” (Fig. 3a-iv).

Figure 3b highlights how only 53.9±2.6% (mean ± SEM) of the origami binding sites detected in conventional DNA-PAINT during the pre-wash direct phase appeared during the post-wash direct phase. Fig. S4 shows that such site loss happens progressively over the pre-wash imaging phase and is further enhanced by the washing steps. Progressive site loss is expected for DNA-PAINT and has been previously documented and ascribed to photoinduced chemical damage of the docking motifs.^27^ Shear forces exerted while washing might account for additional origami damage.

Fig. 3c shows a fraction of retained sites fR= 52.2±1.8% for PD-PAINT on tiles with *d*=5 nm, as measured between the direct phase and the PD phase. The nearly identical values of f_R_ found for PD-PAINT and the control experiment prove that essentially all sites that would remain functional over the course of a comparable DNA-PAINT experiment will produce a PD-PAINT signal for tiles with *d*=5 nm.

The fraction of retained sites however decreases with increasing tethering distance, to 27.7±1.1% for *d*=10 nm and 18.3±0.9% for *d*=15 nm (Fig. 3d,e). Such an additional decrease in viable sites is unexpected, as S1 and S2 were predicted to have similar dimerization probabilities within the tested *d*-range. The imperfect addressability of overhangs on the tile, first reported by Strauss et al.,^28^ and assessed for our tiles in Fig. S1, might be partially responsible for the *d*-dependent site loss, given the change in the position of D2 between the three different designs (Fig. 3c-e). However, the trend we detected does not correlate with the published addressability maps.^28^

Another possible explanation for the observed *d*-dependence of f_R_ is a difference in S1-S2 dimerization kinetics for different tethering distances. While the free energy calculations in Fig. 2 indicate that the equilibrium dimerization probability should be independent of *d* within the studied range, they do not offer direct insight into the kinetics of the process. Long dimerization times, increasing with *d*, could result in a distance-dependent reduction in the fraction of S1 sites becoming available over the finite PD phase.

To gather detailed experimental insights into the kinetics and thermodynamics of PD-PAINT we carried out a single molecule statistical analysis of the “blinking” behavior of individual origami sites. Fig. 4a shows the normalized cumulative distribution function (CDF) of the dark times, which are defined as the time intervals between subsequent blinks. Data are shown for PD-PAINT experiments at all tethering distances, comparing direct and PD phases, and as a control also for conventional DNA-PAINT, comparing both pre- and post-wash direct phases.

**Figure 4.**
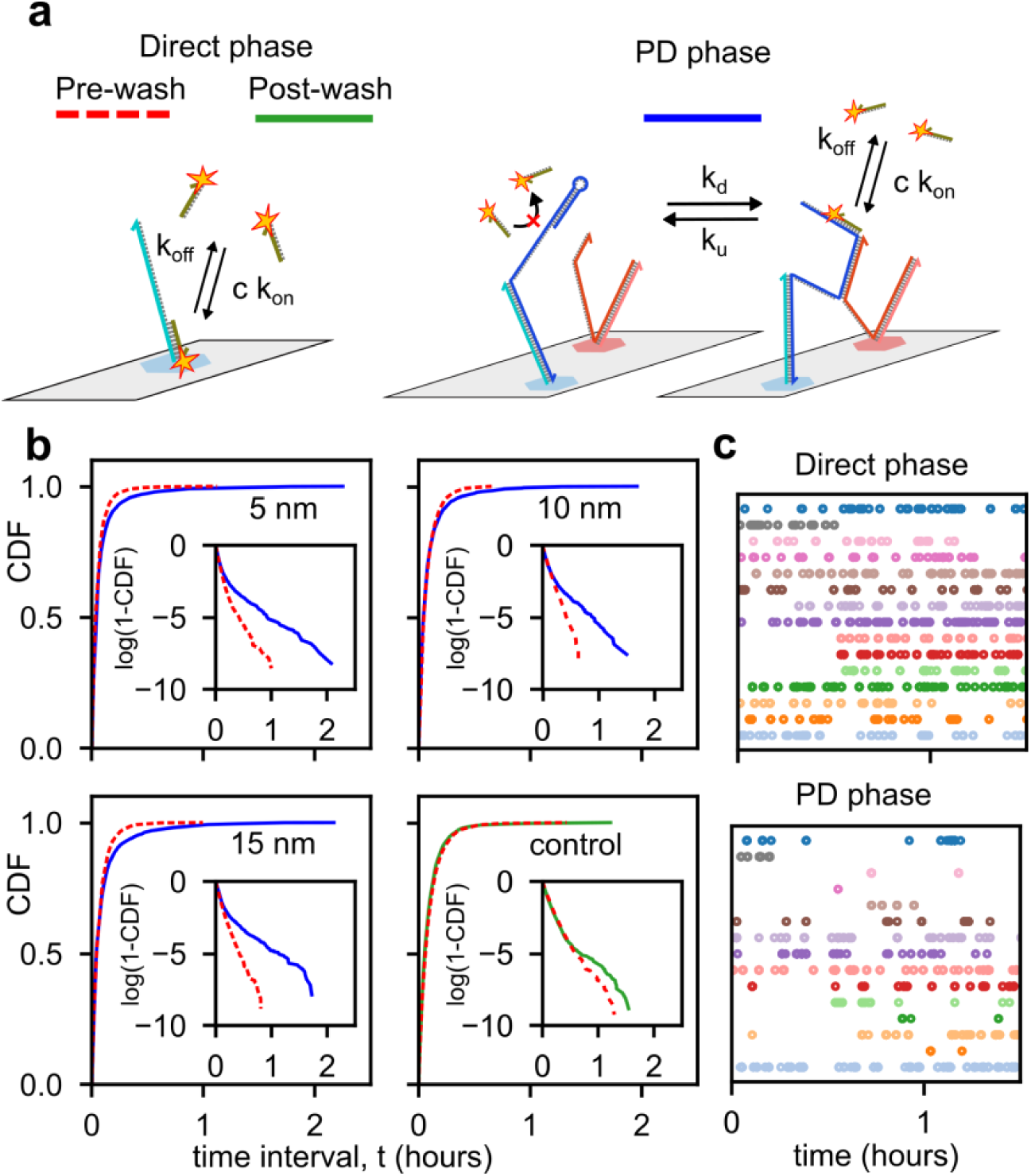
Single-molecule kinetic and thermodynamic assessment of PD-PAINT. **a:** Kinetic processes involved in conventional DNA-PAINT, as carried out in the direct imaging phases, and in PD-PAINT (PD-phases). In the latter situation, reversible dimerization and unbinding of S1 and S2 takes place, resulting in extended dark periods in which the P1 binding site is rendered inaccessible. **b:** Cumulative Distribution Function (CDF) of the dark times recorded in (pre- and post-wash) direct phases and PD phases (legend in panel **a**) for PD-PAINT experiments with different S1-S2 separation *d* and a control DNA-PAINT experiment. Note the greater prominence of long dark times in PD phases as opposed to direct phases. Insets: logarithm of (1-CDF) displaying an exponential trend for data collected in the direct phases, and the substantial deviation observed in PD phases. See SI section 1.15, 1.16, Fig. S2 and Table S3, S4 and S5 for analytical and numerical analysis of the dark time distributions and the estimation of the rate constants in panel **a. c:** Examples of blinking sequences recorded in the PD (top) and direct (bottom) phases, where each symbol represents an event, and gaps between them the dark times. Extended dark periods are visibly more common in PD-PAINT data.

For all direct imaging phases, in PD-PAINT and control experiments (both pre- and post-wash), the distributions closely follow an exponential trend, visible in the insets of Fig. 4b. This functional form is expected for DNA-PAINT experiments, and results from the second-order kinetics of the imager binding to the docking domain with rate constant k_on_. The latter is estimated for all direct experimental phases by fitting the respective dark time distributions, and found in good agreement with previous observations for the P1 imager binding to its docking site^29^ (k_on_ = 1.48 − 2.50 × 10^6^ *M*^−1^ *s*^−1^ using [P1] = 2 nM, see SI section 1.16, Fig. S2 and Table S3). For all PD phases, in turn, the distributions display a clear deviation from the corresponding direct-phase curves, with a substantial excess of long dark times (Fig. 4b and insets). Direct inspection of the sampled blinking sequences, exemplified in Fig. 4c (bottom), reveal that the excess of long dark times follows from an intermittent behavior of PD-PAINT sites, which display streaks of frequent bright events interrupted by long periods of inactivity. By contrast, a more regular imager attachment rate is observed in direct phases (Fig. 4c top).

We ascribe the intermittent blinking of PD-PAINT sites to the reversibility of S1-S2 dimerization, which leads to transitions between the dimerized state, in which the S1 docking site is accessible to P1, and the un-dimerized state, which renders the docking site inaccessible (Fig. 4c). We label the (first-order) rate constants of S1-S2 dimerization and unbinding as k_d_ and k_u_, and demonstrate that by allowing for this reversibility the CDF of PD-PAINT dark times can be described by a hyper-exponential distribution, which we can fit to the data to extract values for the two rate constants (SI section 1.16, Fig. S2 and Table S3, S4 and S5). To overcome limitations of the analytical description, which fails to capture the finite sampling time of the experimental datasets, we also carry out explicit stochastic simulations of the blinking pattern, modeled as a continuous Markov process. Optimization of the simulated dark time distributions allows for estimating k_d_ and k_u_, which are in good agreement with the outcomes of the analytical fit (SI section 1.16, Fig. S2 and Table S4).

The timescales associated to the extracted kinetic rates, 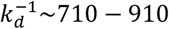 s, 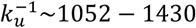 s and (k_on_ [*P*1])^−1^ ∼180 − 200 s, are in all cases substantially shorter than the duration of the PD imaging phase (1 × 10^4^ − 1.3 × 10^4^s). Therefore, the ensemble of detectable S1-S2 pairs should comfortably reach equilibrium within the experimental timeframe.

The equilibration kinetics was also directly probed by measuring the first passage time, defined as the time interval between the addition of the removal strand R and the detection of the first bright event on each S1-S2 site. Results, summarized in Fig. S3 confirm that the first passage times are, for all tested *d*-values, substantially shorter compared to the duration of the PD phase. Consequently, all origami sites in which both S1 and S2 are present and chemically intact should produce blinks within the observation timescales, and kinetics can be ruled out as the cause of the excess site loss measured for *d*=10 nm and 15 nm.

The consistency between the blinking kinetics of the detectable PD-PAINT sites and the theoretical models (both analytical and numerical) indicates that also the excess site loss observed for *d*=10 nm and 15 nm is likely to derive from non-ideal features in a sub-population of sites, rather than from intrinsic design issues with the PD-PAINT probes. From the experimental values of k_d_ and k_u_ we can extract the probabilities of S1-S2 dimerization, and compare it with the results of simulations in Fig. 2b. The latter are found to overestimate the stability of the S1-S2 dimer, a result which might derive from limitations of the coarse-grained description of DNA we adopt, or from necessary simplifications of the simulated system where, for instance, the origami is not explicitly included (see Supporting Information). We point out that the lower S1-S2 dimerization probability detected in experiments has little impact on the performance of PD-PAINT, having demonstrated through our single-molecule analysis that the kinetics of S1-S2 binding and unbinding is sufficiently fast to guarantee that bright events are recorded within typical experimental time-scales whenever the two probes are within reach – which is ultimately the key requirement.

The DNA origami platform also enabled several control experiments to confirm that the number of false positives, i.e. signals detected from S1 when no S2 structures are in proximity, is low and that shielding of S2 worked as expected. In an experiment where only S1 was present, 1.89% of the origami binding sites imaged during the direct phase were detectable in the PD phase, indicating a very low false-positive rate (Fig. S6e). Similarly, protection of S2 with the shield B avoided interaction with S1 when applied at a concentration used during attachment (Fig. S6c) whereas unprotected S2 is found to dimerize to tethered S1 strands (Fig. S6b). Finally, we confirmed that adding the shield remover R does not affect closed S1 loops in the absence of S2 (Fig. S6d).

Having assessed the performance of PD-PAINT in-silico and on test samples, we applied it to the detection of proximity between two proteins in a biological sample (Fig. 5). In particular, we targeted the cardiac ryanodine receptor (RyR) and the junctional protein junctophilin (JPH)^30,31^ in fixed cardiac cells. These proteins had been previously shown to localize in close proximity, with a subset of JPH found within ≤50 nm of RyRs.

**Figure 5.**
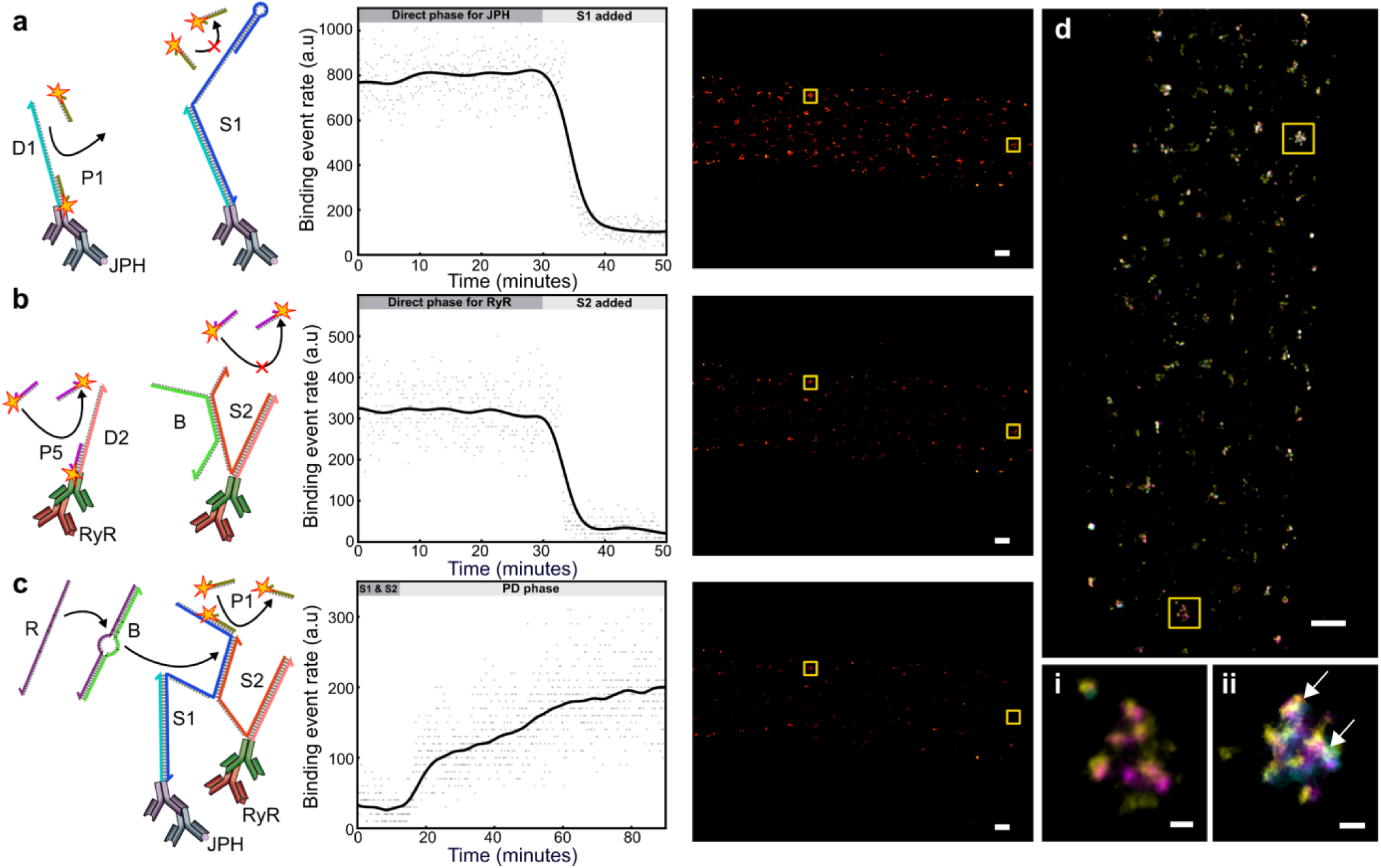
PD-PAINT assay of the proximity of cardiac proteins JPH and RyR in isolated cardiomyocytes. Shown in a-c from left to right: DNA process schematic (not to scale), single molecule time trace of events, and final rendered image. **a:** Initial imaging of P1 docking sites. S1 introduced at ∼500 nM binding to D1 and preventing further P1 imager docking events. Single molecule events are reduced to background levels within an approximate 10 min incubation period. **b:** Following an Exchange-PAINT protocol P5 imagers are then used to probe RyR on D2. Pre-annealed S2-B then bind to D2 preventing further RyR sampling, taking ∼10 minutes to reduce events. **c:** The addition of the displacement sequence R removes B and enables S1-S2 dimerization when in close enough proximity. This process is stochastic and the single molecule events gradually increase over time as more S1 loops are opened. **d:** Overlay of super-resolution images obtained at each stage (a-c, rotated 90°) showing JPH (yellow), RyR (magenta), and PD-PAINT signal (Cyan). Magnification of boxed regions left (i) where JPH and RyR signal is clearly present within the cluster but are seemingly not in close molecular proximity for S1-S2 dimerization and (ii) showing all 3 signals present (arrows highlight strongly overlapping signals which appear white). Frame numbers used to generate super-resolution images: D1 & D2 ∼30k frames & open S1 sites ∼60k frames. Scale bars: (a-d) 1µm, (i-ii) 100 nm.

The two epitopes were first targeted by two distinct primary antibodies (ABs). Secondary ABs conjugated to the D1 and D2 strands (see Fig. S7) were then applied, targeting the primary ABs for JPH and RyR, respectively.

Prior to adding S1, the JPH epitopes were directly imaged using the docking site on D1 with imager P1, as shown in Fig. 5a. A clear signal and the distinct patterns expected from these proteins were found.^32^ Upon addition of S1, blocking of the P1 docking site on D1 resulted in a rapid drop in event rate to background levels, just as observed with the origami platform. Following an Exchange-PAINT step^24^ to wash out P1 imagers, we then directly imaged RyRs using newly added P5 imagers, complementary to a domain on the D2 strands. We detected a broadly similar pattern to the one found for JPH (Fig, 5b), as previously observed in dual-color super-resolution studies of RyR-JPH distribution.^31^ As expected, the addition of S2 and B, capping D2, rapidly suppressed the P5 imager signal indicating successful attachment of S2-B.

Finally, upon re-introducing P1 and adding the remover strand R, bright events resumed following the dimerization of S1 and S2, as expected, given the proximity of some RyR and JPH epitopes^4^ (Fig. 5c). The rate of events increases over time, with a transient of a few minutes, in line with the dimerization kinetics observed for the origami platform.

Fig. 5d demonstrates that although JPH and RyR often appear co-localized within the same molecular cluster, the PD-PAINT signal is not always present, indicating that in some cases the JPH-RyR proximity is not sufficient to allow direct interaction and S1-S2 dimerization. We note that false positives resulting from conventional co-localization approaches are expected where the local overlap of signal in different channels is often artifactual, arising simply from limited spatial resolution in the more complex biological environment. The improved PD-PAINT assay avoids this reliance on spatial resolution and instead employs a physical distance criterion that is independent of imaging resolution.

Note that JPH appears as the most abundant protein (Fig. 5a), RyR is slightly less abundant (Fig. 5b), while the weakest signals are detected for RyR-JPH pairs with PD-PAINT. This is consistent with previous observations on RyRs and JPH.^4^ The number of RyR-JPH pairs are bounded by the smaller number of RyRs and the lesser abundance of PD signal pairs is therefore consistent with expectations.

A second pair of cardiac proteins, RyRs and the intrinsic sarcoplasmic reticulum membrane protein SERCA^33^, were imaged in a similar way with PD-PAINT in cardiac tissue sections, which provide an environment very challenging for optical superresolution imaging. The experiment yielded comparable results to the observations above, as shown in Fig. S8.

The dependency of PD-PAINT signals on the molecular proximity of S1 and S2 was further demonstrated in an experiment that serves as a negative control (Fig. S9). For this experiment, collagen type VI (ColVI) and RyR were labeled with primary and secondary antibodies to carry D1/S1 and D2/S2 constructs, respectively (Fig. S9a). RyR and ColVI have a broadly similar distribution pattern in cardiomyocytes but are separated by the cell membrane and thus not in direct molecular contact. When the two antibody populations were individually imaged using conventional DNA-PAINT targeted towards the D1 and D2 docking strands (Fig. S9b,c), the resulting two-color DNA-PAINT image exhibited overlap in some areas, an occurrence that would be interpreted as molecular co-localization (Fig. S9d). When the same areas were imaged using PD-PAINT, after S1 and S2 were tethered to D1 and D2, no signal was detected above background levels (Fig. S9e). This suggests that the areas of overlap observed in the two-color DNA-PAINT image are false positives and that no epitopes of ColVI and RyR are located within close proximity to each other. This is consistent with the expected distribution of collagen and RyRs - ColVI and RyR would be expected to be separated by 40 nm or more based on myocyte ultrastructure determined by electron microscopy.^34^ Note that the probability of S1-S2 dimerization predicted by MC simulations at such separations is, consistently, negligibly small (Fig. 2b), even if taking into account a possibly reduced distance of the proximity probes due to the spatial extent of antibody labels (which can account for up to ∼20 nm).

One of the key advantages of DNA-PAINT is that of enabling the quantification of the number of accessible docking sites, and hence of the labeled epitopes, through the qPAINT method.^23^ Quantitative estimates of protein-pair numbers are critical, for example, in assays that test for changes in protein-protein signaling in response to experimental interventions. Some types of proximity detection have been shown to be prone to saturation which can preclude detecting changes in such experiments.^35^

We demonstrate that qPAINT analysis can be seamlessly integrated with PD-PAINT, just like with regular DNA-PAINT. The qPAINT approach simply relies on recording dark-time distributions from areas with an unknown number of docking sites (*N)*, and then exploiting the linear relationship between the inverse mean dark time and *N*. A calibration performed on data with known number of docking sites then enables the extraction of *N* from the measured mean dark time. Using the experimental data collected on origami test samples we demonstrate that, regardless of the differences in the distributions of dark times between PD-PAINT and conventional DNA-PAINT (discussed above), the linear relationship between the inverse mean dark time and *N* persists, proving the applicability of the technique (see Supporting Information and Fig. S10).

In principle, care must be taken when applying the qPAINT analysis to PD-PAINT data collected from samples with non-uniform S1-S2 separation, as would be common in biological scenarios. Indeed, a dependence of the blinking kinetics on *d* could bias the results if one relied on calibration data acquired at uniform *d*. Comparison between qPAINT calibration curves recorded on origami data for *d=*5 nm, 10 nm and 15 nm reveals relatively similar slopes, proving that at least within this range the bias should be limited (Fig. S10 and Table S1). It is however likely that the most precise qPAINT estimates in biological samples can be obtained in cases where the distance between target epitopes is uniform throughout the sample or approximately known a priori, e.g. for proteins clustering in known morphologies. A possible distance dependent site loss, similar to the one we noticed with origami, could introduce additional bias, along with other experimental artifacts common in all DNA-PAINT measurements, such as imperfect epitope labeling and photo damage of the docking strands.

## Conclusion

In summary, we present Proximity-Dependent PAINT, a technique which allows imaging of the distribution of protein pairs, or other biological targets in close proximity, with nanoscale resolution. As opposed to conventional multi-channel-imaging co-localization techniques, proximity detection is decoupled from the imaging resolution, which is especially important in biological cell and tissue samples where the achievable resolution can vary greatly due to refractive index inhomogeneities and related optical challenges. Indeed, with PD-PAINT the proximity range can be smaller than the imaging resolution, a distinct advantage that the proposed method shares with FRET imaging. However, in contrast to FRET, PD-PAINT does not require specific dye pairs, eliminates the need for a more complex excitation and detection setup, and does not suffer from the effects of specific molecular orientations which can complicate the interpretation of FRET experiments. In addition, PD-PAINT can be efficiently multiplexed, a more challenging task with FRET-based approaches.

The viability of PD-PAINT is assessed through coarse-grained molecular simulations and experiments on DNA origami test tiles, the latter enabling single-molecule analysis of the kinetic and thermodynamic features of the platform.

We show that PD-PAINT is compatible with the complexities of biological environments by imaging different epitopes of primary antibodies and different epitopes of the ryanodine receptor, a large protein, in fixed cardiac tissue sections. Owing to the modularity of the DNA nanostructures used, PD-PAINT can be implemented relatively easily and with little penalty as an additional step in any Exchange-PAINT experiment.^24^ With modern sensitive and affordable cameras (Exchange-) PD-PAINT is straightforward to implement on a conventional fluorescence microscope, the equipment required can be assembled from components with a relatively modest budget.^36^

We demonstrated the application of PD-PAINT to cellular preparations using a secondary antibody system which due to its size could extend the capture radius of the PD-PAINT system. In the applications shown here this has only a minor effect since one of the protein partners, the RyR, is itself very large, with a diameter of ∼30 nm. For potentially smaller protein pairs, PD-PAINT can be readily adapted by conjugating the proximity-detection strands S1 and S2 to compact markers such as nanobodies, aptamers and other emerging probes.^37,38^

PD-PAINT is fully compatible with multi-color imaging and can be implemented in addition to conventional DNA-PAINT of multiple other channels, essentially without performance penalty. This is demonstrated here by following an Exchange-PAINT protocol,^24^ to image the protein complex as well as the individual protein epitopes in separate channels (Fig. S12).

Finally, we show that PD-PAINT can be seamlessly combined with qPAINT analysis for measuring emitter numbers.^23^ Guided by simulations and single-molecule experimental analysis, one could foresee design changes to the basic PD-PAINT machinery aimed at optimizing its performance in specific settings. For example, one could fine tune the range of proximity-detectability. The PD-PAINT system could also be modified to sense the proximity of more than two targets, similar to what has been previously shown for PLA.^39^ Using the vast combinatorial space of DNA-sequence design, it should be possible to design orthogonal imaging probes that analogously to S1 and S2 expose different docking strands when in close proximity. Based on this principle, a virtually unlimited number of distinct protein pairs can be detected within a single sample.

Increasing the affinity between the imager and the S1 docking domain, e.g. by adjusting the CG/AT ratio or by extending the number of bases in the stem loop, in combination with an adjusted imager concentration, would enable a near-permanent labeling of open S1-loop structures with single imagers. In this configuration, the PD-PAINT machinery can be exploited to render other super-resolution imaging modalities proximity sensitive, such as STED, SIM, (d)STORM, (f)PALM, or even widefield and confocal microscopy.^13,14,16,17,19^

## ASSOCIATED CONTENT

### Supporting Information

Materials and experimental methods, simulation methods, a table showing qPAINT inverse mean dark times per emitter, a table of the mean first passage times for bright events to be regenerated in PD-PAINT for different distances, a table with estimates of the rates of imager hybridization, a table showing the analytical estimates of the rates of the processes underlying the blinking kinetics of PD-PAINT sites, a table with numerical estimates of the rate constants using a Hidden Markov Model, tables showing DNA sequences, a table showing expected binding timescales for the dissociation of the P1 imager to the docking site on S1, a table with the windows used in the simulations, a figure showing the validation of the accessibility of D1 & D2 overhangs on DNA origami, a figure of the analytical and numerical fits of the dark-time cumulative distribution functions, a figure showing the analysis of event first passage time for PD-PAINT, a figure evaluating site loss during conventional DNA-PAINT, a figure showing an example event-rate time trace on origami, a figure showing several PD-PAINT control experiments, a figure showing spectra of DNA conjugation products,, a figure showing an additional PD-PAINT investigation of a protein pair, a figure showing a negative control experiment for PD-PAINT, a figure showing qPAINT analysis, a figure showing gels and AFM images of DNA origami synthesis and a figure showing multiplexed PD-PAINT ratio-imaging. This material is available in the Supporting Information document found online.

## AUTHOR INFORMATION

### Author Contributions

The manuscript was written through contributions of all authors.

### Funding Sources

This work was supported by the Human Frontier Science Program (No. 0027/2013) and the Engineering and Physical Sciences Research Council of the UK (No. EP/N008235/1) and Biotechnology and Biological Sciences Research Council Grants BB/P026508/1 and BB/T007176/1. L.D.M. acknowledges funding from a Royal Society University Research Fellowship (UF160152) and from the European Research Council (ERC) under the Horizon 2020 Research and Innovation Programme (ERC-STG No 851667 NANOCELL). W.T.K. acknowledges support from an EPSRC DTP studentship.

### Conflict of Interest Disclosure

T.L., W.T.K., A.H.C., L.D.M. and C.S. are inventors on patent application GB1904095.5.

## ACKNOWLEDGMENT

We thank Dr Ruisheng Lin for assistance with the imaging setup. We gratefully acknowledge helpful discussions with Prof. Ralf Jungmann on DNA origami design and synthesis.

## Notes

### Competing Interest Statement

The authors have declared no competing interest.

## REFERENCES

(1) Jones, S.; Thornton, J. M. Principles of Protein-Protein Interactions. Proc. Natl. Acad. Sci. 1996, 93 (1), 13–20. https://doi.org/10.1073/pnas.93.1.13.

(2) Phizicky, E. M.; Fields, S. Protein-Protein Interactions: Methods for Detection and Analysis. Microbiol. Rev. 1995, 59 (1), 94–123. https://doi.org/10.1128/mmbr.59.1.94-123.1995.

(3) Fields, S.; Song, O. K. A Novel Genetic System to Detect Protein-Protein Interactions. Nature 1989, 340 (6230), 245–246. https://doi.org/10.1038/340245a0.

(4) Jayasinghe, I.; Clowsley, A. H.; Lin, R.; Baddeley, D.; Michele, L. Di; Soeller, C.; Jayasinghe, I.; Clowsley, A. H.; Lin, R.; Lutz, T.; Harrison, C.; Green, E.; Baddeley, D.; Michele, L. Di; Soeller, C. True Molecular Scale Visualization of Variable Clustering Properties of Ryanodine Receptors. Cell Rep. 2018, 22 (2), 557–567. https://doi.org/10.1016/j.celrep.2017.12.045.

(5) Söderberg, O.; Gullberg, M.; Jarvius, M.; Ridderstråle, K.; Leuchowius, K. J.; Jarvius, J.; Wester, K.; Hydbring, P.; Bahram, F.; Larsson, L. G.; Landegren, U. Direct Observation of Individual Endogenous Protein Complexes in Situ by Proximity Ligation. Nat. Methods 2006, 3 (12), 995–1000. https://doi.org/10.1038/nmeth947.

(6) Fredriksson, S.; Gullberg, M.; Jarvius, J.; Olsson, C.; Pietras, K.; Gústafsdóttir, S. M.; Östman, A.; Landegren, U. Protein Detection Using Proximity-Dependent DNA Ligation Assays. Nat. Biotechnol. 2002, 20 (5), 473–477.

(7) Gullberg, M.; Gústafsdóttir, S. M.; Schallmeiner, E.; Jarvius, J.; Bjarnegård, M.; Betsholtz, C.; Landegren, U.; Fredriksson, S. Cytokine Detection by Antibody-Based Proximity Ligation. Proc. Natl. Acad. Sci. U. S. A. 2004, 101 (22), 8420–8424. https://doi.org/10.1073/pnas.0400552101.

(8) Söderberg, O.; Leuchowius, K. J.; Gullberg, M.; Jarvius, M.; Weibrecht, I.; Larsson, L. G.; Landegren, U. Characterizing Proteins and Their Interactions in Cells and Tissues Using the in Situ Proximity Ligation Assay. Methods 2008, 45 (3), 227–232. https://doi.org/10.1016/j.ymeth.2008.06.014.

(9) Dirks, R. M.; Pierce, N. A. Triggered Amplification by Hybridization Chain Reaction. Proc. Natl. Acad. Sci. U. S. A. 2004, 101 (43), 15275–15278. https://doi.org/10.1073/pnas.0407024101.

(10) Koos, B.; Cane, G.; Grannas, K.; Löf, L.; Arngården, L.; Heldin, J.; Clausson, C. M.; Klaesson, A.; Hirvonen, M. K.; De Oliveira, F. M. S.; Talibov, V. O.; Pham, N. T.; Auer, M.; Danielson, U. H.; Haybaeck, J.; Kamali-Moghaddam, M.; Söderberg, O. Proximity-Dependent Initiation of Hybridization Chain Reaction. Nat. Commun. 2015, 6. https://doi.org/10.1038/ncomms8294.

(11) Koos, B.; Andersson, L.; Clausson, C.-M.; Grannas, K.; Klaesson, A.; Cane, G.; Söderberg, O. Analysis of Protein Interactions in Situ by Proximity Ligation Assays. In High-Dimensional Single Cell Analysis: Mass Cytometry, Multi-parametric Flow Cytometry and Bioinformatic Techniques; Fienberg, H. G., Nolan, G. P., Eds.; Current Topics in Microbiology and Immunology; Springer Berlin Heidelberg: Berlin, Heidelberg, 2014; pp 111–126.

(12) Truong, K.; Ikura, M. The Use of FRET Imaging Microscopy to Detect Protein-Protein Interactions and Protein Conformational Changes in Vivo. Curr. Opin. Struct. Biol. 2001, 11 (5), 573–578. https://doi.org/10.1016/S0959-440X(00)00249-9.

(13) Bailey, B.; Farkas, D. L.; Taylor, D. L.; Lanni, F. Enhancement of Axial Resolution in Fluorescence Microscopy by Standing-Wave Excitation. Nature 1993, 366 (6450), 44–48.

(14) Hell, S. W.; Wichmann, J. Breaking the Diffraction Resolution Limit by Stimulated Emission : Stimulated-Emission-Depletion Fluorescence Microscopy. Opt. Lett. 1994, 19 (11), 780–782.

(15) Klar, T. A.; Jakobs, S.; Dyba, M.; Egner, A.; Hell, S. W. Fluorescence Microscopy with Diffraction Resolution Barrier Broken by Stimulated Emission. Proc. Natl. Acad. Sci. 2000, 97 (15), 8206–8210. https://doi.org/10.1073/pnas.97.15.8206.

(16) Betzig, E.; Patterson, G. H.; Sougrat, R.; Lindwasser, O. W.; Olenych, S.; Bonifacino, J. S.; Davidson, M. W.; Lippincott-Schwartz, J.; Hess, H. F. Imaging Intracellular Fluorescent Proteins at Nanometer Resolution. Science 2006, 313 (5793), 1642–1645. https://doi.org/10.1126/science.1127344.

(17) Hess, S. T.; Girirajan, T. P. K.; Mason, M. D. Ultra-High Resolution Imaging by Fluorescence Photoactivation Localization Microscopy. Biophys. J. 2006, 91 (11), 4258–4272. https://doi.org/10.1529/biophysj.106.091116.

(18) Rust, M. J.; Bates, M.; Zhuang, X. W. Sub-Diffraction-Limit Imaging by Stochastic Optical Reconstruction Microscopy (STORM). Nat Methods 2006, 3 (10), 793–795. https://doi.org/Doi10.1038/Nmeth929.

(19) Heilemann, M.; Linde, S. Van De; Schüttpelz, M.; Kasper, R.; Seefeldt, B.; Mukherjee, A.; Tinnefeld, P.; Sauer, M. Subdiffraction-Resolution Fluorescence Imaging with Conventional Fluorescent Probes. Angew. Chemie Int. Ed. 2008, 47, 6172–6176. https://doi.org/10.1002/anie.200802376.

(20) Schnitzbauer, J.; Strauss, M. T.; Schlichthaerle, T.; Schueder, F.; Jungmann, R. Super-Resolution Microscopy with DNA-PAINT. Nat. Protoc. 2017, 12 (6), 1198–1228. https://doi.org/10.1038/nprot.2017.024.

(21) Xu, K.; Babcock, H. P.; Zhuang, X. Dual-Objective STORM Reveals Three-Dimensional Filament Organization in the Actin Cytoskeleton. Nat. Methods 2012, 9 (2), 185–188. https://doi.org/10.1038/nmeth.1841.

(22) Dai, M.; Jungmann, R.; Yin, P. Optical Imaging of Individual Biomolecules in Densely Packed Clusters. Nat. Nanotechnol. 2016, 11 (9), 798–807.

(23) Jungmann, R.; Avendaño, M. S.; Dai, M.; Woehrstein, J. B.; Agasti, S. S.; Feiger, Z.; Rodal, A.; Yin, P. Quantitative Super-Resolution Imaging with QPAINT. Nat. Methods 2016, 13 (5), 439–442. https://doi.org/10.1038/nmeth.3804.

(24) Jungmann, R.; Avendaño, M. S.; Woehrstein, J. B.; Dai, M.; Shih, W. M.; Yin, P. Multiplexed 3D Cellular Super-Resolution Imaging with DNA-PAINT and Exchange-PAINT. Nat. Methods 2014, 11 (3), 313–318. https://doi.org/10.1038/nmeth.2835.

(25) Ouldridge, T. E.; Louis, A. A.; Doye, J. P. K. DNA Nanotweezers Studied with a Coarse-Grained Model of DNA. Phys. Rev. Lett. 2010, 104 (17), 1–4. https://doi.org/10.1103/PhysRevLett.104.178101.

(26) Ouldridge, T. E.; Louis, A. A.; Doye, J. P. K. Structural, Mechanical, and Thermodynamic Properties of a Coarse-Grained DNA Model. J. Chem. Phys. 2011, 134 (8). https://doi.org/10.1063/1.3552946.

(27) Blumhardt, P.; Stein, J.; Mücksch, J.; Stehr, F.; Bauer, J.; Jungmann, R.; Schwille, P. Photo-Induced Depletion of Binding Sites in DNA-Paint Microscopy. Molecules 2018, 23 (12), 1–27. https://doi.org/10.3390/molecules23123165.

(28) Strauss, M. T.; Schueder, F.; Haas, D.; Nickels, P. C.; Jungmann, R. Quantifying Absolute Addressability in DNA Origami with Molecular Resolution. Nat. Commun. 2018, 9 (1), 1–7. https://doi.org/10.1038/s41467-018-04031-z.

(29) Jungmann, R.; Steinhauer, C.; Scheible, M.; Kuzyk, A.; Tinnefeld, P.; Simmel, F. C. Single-Molecule Kinetics and Super-Resolution Microscopy by Fluorescence Imaging of Transient Binding on DNA Origami. Nanoletters 2010, 10, 4756–4761. https://doi.org/10.1021/nl103427w.

(30) Jayasinghe, I. .; Baddeley, D.; Kong, C. .; Wehrens, X. H. .; Cannell, M. .; Soeller, C. Nanoscale Organization of Junctophilin-2 and Ryanodine Receptors within Peripheral Couplings of Rat Ventricular Cardiomyocytes. Biophys. J. 2012, 102 (5), L19–21.

(31) Munro, M.; Jayasinghe, I. D.; Wang, Q.; Quick, A.; Baddeley, D.; Wehrens, X. H. .; Soeller, C. Junctophilin-2 in the Nanoscale Organisation and Functional Signalling of Ryanodine Receptor Clusters in Cardiomyocytes. J. Cell Sci. 2016, 129 (23), 4388–4398.

(32) Munro, M. L.; Shen, X.; Ward, M.; Ruygrok, P. N.; Crossman, D. J.; Soeller, C. Highly Variable Contractile Performance Correlates with Myocyte Content in Trabeculae from Failing Human Hearts. Sci. Rep. 2018, 8 (1), 1–13. https://doi.org/10.1038/s41598-018-21199-y.

(33) MacLennan, D.; Kranias, E. Phospholamban: A Crucial Regulator of Cardiac Contractility. Nat. Rev. Mol. Cell Biol. 2003, 4 (7), 566–577.

(34) Morita, T.; Shimada, T.; Kitamura, H.; Nakamura, M. Demonstration of Connective Tissue Sheaths Surrounding Working Myocardial Cells and Purkinje Cells of the Sheep Moderator Band. Arch. Histol. Cytol. 1991, 54 (5), 539–550. https://doi.org/10.1679/aohc.54.539.

(35) Mocanu, M. M.; Váradi, T.; Szöllosi, J.; Nagy, P. Comparative Analysis of Fluorescence Resonance Energy Transfer (FRET) and Proximity Ligation Assay (PLA). Proteomics 2011, 11 (10), 2063–2070. https://doi.org/10.1002/pmic.201100028.

(36) Ma, H.; Fu, R.; Xu, J.; Liu, Y. A Simple and Cost-Effective Setup for Super-Resolution Localization Microscopy. Sci. Rep. 2017, 7 (1), 1–9. https://doi.org/10.1038/s41598-017-01606-6.

(37) Schlichthaerle, T.; Eklund, A. S.; Schueder, F.; Strauss, M. T.; Tiede, C.; Curd, A.; Ries, J.; Peckham, M.; Tomlinson, D. C.; Jungmann, R. Site-Specific Labeling of Affimers for DNA-PAINT Microscopy. Angew. Chemie - Int. Ed. 2018, 57 (34), 11060–11063. https://doi.org/10.1002/anie.201804020.

(38) Sograte-Idrissi, S.; Oleksiievets, N.; Isbaner, S.; Eggert-Martinez, M.; Enderlein, J.; Tsukanov, R.; Opazo, F. Nanobody Detection of Standard Fluorescent Proteins Enables Multi-Target DNA-PAINT with High Resolution and Minimal Displacement Errors. Cells 2019, 8 (1), 48. https://doi.org/10.3390/cells8010048.

(39) Schallmeiner, E.; Oksanen, E.; Ericsson, O.; Spångberg, L.; Eriksson, S.; Stenman, U. H.; Pettersson, K.; Landegren, U. Sensitive Protein Detection via Triple-Binder Proximity Ligation Assays. Nat. Methods 2007, 4 (2), 135–137. https://doi.org/10.1038/nmeth974.

