## Supporting Information for "Detecting nanoscale distribution of protein pairs by proximity dependent super-resolution microscopy"

### 1 Materials and Experimental Methods

#### 1.1 DNA Origami production

All oligonucleotides (staples) used in the construction of the origami were purchased from Integrated DNA Technologies (Coralville, Iowa, United States), purified by standard desalting, prediluted at 100 µM concentration in Tris EDTA buffer pH 8.0, and shipped in 96 well plates. Oligonucleotides were stored frozen at -20ºC. The single stranded circular viral template (scaffold), M13mp18, was purchased from New England Biolabs (Ipswitch, Massachusetts, United States).
Rothemund Rectangular Origami (RRO) with various 3' overhangs were manufactured as detailed in a previous publication.[1] Briefly, the Python program *Picasso* was used to generate staple sequences which fold to yield an RRO with 3' overhangs to various specified locations on a single face of the planar origami. On the reverse face, eight DNA strands were given 5' biotin modifications for anchoring.
Sequences of DNA oligonucleotides without end modifications are listed in Table S8. There are four sets of overhangs for 3 different origami structures that differ in the distance between D1 and D2 overhangs: D1 (red), D2a (blue), D2b (green), D2c (pink). Origami with specific combinations of overhangs are made by substituting the overhang bearing strands for the corresponding strands lacking overhangs. The sequences of the eight 5'-biotinylated staples, present in all designs, are listed in Table S7.
RROs were manufactured at a total volume of 40 µL in 1x Tris EDTA buffer (TE, 10 mM Tris-HCl, 1 mM disodium EDTA, pH 8.0, Sigma) supplemented with 12.5 mM $MgCl_2$ (Magnesium chloride hexahydrate, BioXtra,> 99%) such that the final concentration of components was 10 nM scaffold, 10 nM biotinylated oligos, 100 nM oligos without 3' overhangs, 1 µM oligos with 3' overhangs. Assembly of RROs was facilitated by thermal annealing using a Techne TC-512 thermocycler. The mixture was brought to 80ºC to eliminate secondary structure, followed by reducing from 60ºC to 40ºC at 3 minutes 12 seconds per degree.

#### 1.2 Agarose Gel

Gels were manufactured at 1.5% agarose (Sigma) in 1x TBE (89 mM Tris, 89 Boric Acid, 2 mM EDTA, Thermo Scientific), supplemented by 10 mM $MgCl_2$. To visualize DNA, the intercalating dye Sybr Safe (SYBR Safe DNA gel stain, 10,000x concentration in DMSO, Invitrogen) was added to the molten gel before casting. A volume of 20 µL of the DNA origami solutions (concentration 10 nM) were mixed with 6x Blue Juice LB loading dye (Invitrogen) and pipetted into each well. Control wells were filled with the origami staples and the (annealed) scaffold. 100 bp ladder (Invitrogen) was added to the first and last wells. 1x TBE was used as a running buffer.
A voltage of 90 V (6 V $cm^{-1}$) was applied to the gel by a Biorad Power Pack Basic for 90 minutes using a 15 cm horizontal gel box (ThermoScientific) in ice.
The agarose gel was visualized using a UV transilluminator (Syngene Dyversity 4), and images captured with a monochrome camera (BFS-U3-63S4M-C USB 3.1 Blackfly S, Monochrome Camera, FLIR Systems) using a 12 mm lens (12mm UC Series Fixed Focal Length Lens, Edmund Optics).

#### 1.3 Atomic Force Microscopy

Correct folding of the 90 nm x 70 nm RRO was confirmed by atomic force microscopy (AFM). After thermal annealing, 1 μL of the DNA origami sample was mixed with 9 μL of 12.5 mM MgCl2 in 1x TE buffer, and pipetted on a mica sheet which had been freshly cleaved using a clean razor blade. The origami sample was incubated with the mica for 10 minutes, followed by two cycles of washing by addition of 300 μL of ultrapure water, and drying with nitrogen for 3 minutes.

AFM was carried out using an MFP-3D Infinity AFM (Asylum Research). BudgetSensors AFM probes (silicon tip, aluminum reflex coating, nominal frequency 300 kHz, stiffness 40 N $m^{-1}$) were used to acquire images in tapping mode, which were analyzed in Gwyddion.[2]

#### 1.4 PD-PAINT materials and sample preparation for imaging

The sequences of the PD-PAINT probes (Table S6) were designed and analyzed with the NUPACK web application (www.nupack.org). Strands were purchased from Eurofins Genomics (Eurofins Scientific, Luxemburg) and IDT (Integrated DNA Technologies, Coralville) purified by high-performance liquid chromatography (HPLC)-purification. Lyophilized DNA for antibody conjugation was resuspended in phosphate buffered saline (PBS, pH 7.4, Sigma-Aldrich), all other DNA was resuspended and stored in Tris-EDTA (TE, pH 8.0, Sigma-Aldrich). The concentration of the reconstituted oligonucleotides was acquired from the DNA absorbance peak (260 nm) and the respective dye absorbance peaks on a NanoDrop spectrophotometer (Thermo Fisher Scientific, Waltham). For biological experiments the extended docking strands D1 and D2 were modified with Cyanine dyes Cy3 and Cy5, respectively, to aid both the quantification of conjugation to secondary antibodies and in identifying regions of interest within biological samples using widefield imaging. Fluorescent emission from these dyes was rapidly photobleached prior to DNA-PAINT imaging and thus did not interfere with super-resolution measurements.

#### 1.5 DNA-antibody conjugation.

Antibody conjugation of extended docking strands D1 and D2 followed a click-chemistry protocol described by Schnitzbauer et al.[1] using AffiniPure Goat Anti-Mouse or Anti-Rabbit IgG (H+L) secondary antibodies (affinity purified and azide-free (115-005-003 & 111-005-003), Jackson ImmunoResearch, PA). For super-resolution imaging, DNA-PAINT buffer (PBS, 500 mM NaCl, pH 8.0) was used, see "buffer C" in a previous publication.[3] Conjugations were quantified by measurements of the dye's absorbance peaks on the modified oligonucleotides, prior to conjugation and afterwards, see Fig. S7. The dye modified oligonucleotide absorbance at the dye peak was scaled to match the absorbance obtained at the same peak for the conjugation. The difference in the absorbance values at 280 nm were attributed to the contribution of the newly conjugated antibody. This was then used to estimate the degree of labeling with typically >1-3 oligonucleotides per antibody being achieved[4] following this procedure:

$$C_{\text{dye}} = A_{\text{dye}}/\varepsilon_{\text{dye}}l \qquad \text{Eq. S 1}$$

$$C_{\text{Ab}} = \left(A_{280} - A_{280\text{ scaled oligo}}\right)/\varepsilon_{\text{Ab}}l \qquad \text{Eq. S 2}$$

$$\text{Ratio}_{\text{oligo/ab}} = C_{\text{dye}}/C_{\text{Ab}}. \qquad \text{Eq. S 3}$$

Here, $C$ is the concentration of either dye or antibody (Ab) worked out by measuring the absorbance, $A$, and using the molar extinction coefficient, $\varepsilon$, and pathlength, $l$.

#### 1.6 Cell and tissue preparation.

Isolated cardiomyocytes were obtained by perfusing 1 mg $mL^{-1}$ Collagenase-II (CLS2, Worthington Biochemical) and 0.1 mg $mL^{-1}$ Protease-I (Sigma) through a cannulated heart using a Langendorff perfusion protocol described previously.[5] Suspended cell solutions were dispensed onto pre-cleaned No.1.5 coverslips treated overnight with 11.9 μg $mL^{-1}$ laminin solution (Life Technologies). After allowing 90 minutes for cell attachment the myocytes were fixed with 2% (w/v) paraformaldehyde (Sigma) in PBS for 10 minutes at room temperature.

Porcine cardiac tissue was fixed in 2% paraformaldehyde (PFA, pH 7.4, Sigma-Aldrich) for 1 hour at 4°C. Tissue samples were cryo-protected using solutions with increasing concentrations of sucrose (up to 30%) and frozen in 2-Methylbutane (Sigma-Aldrich) cooled by liquid nitrogen. Cryo-sections were taken at ~5-15 µm thickness onto pre-cleaned coverslips coated with 0.05% poly-L-lysine (Sigma-Aldrich).

**1.7 PD-PAINT imaging of biological samples**

Coverslips were attached to open-top PMMA imaging chambers (compare Crossman et al.).[6] Biological samples were permeabilized with 0.1% Triton X-100 in PBS for 10 minutes and then incubated with 1% BSA in PBS for 1 hour at room temperature. Primary antibodies, (Ryanodine receptor (RyR) (MA3-916, mouse & HPA016697, rabbit), Junctophilin 2 (JPH2, a custom-made rabbit polyclonal described previously,[7] Serca 2ATPase (SR, ab2861 mouse), and collagen VI (ab6588, rabbit) were incubated overnight at 4°C in an incubation solution containing 1% BSA, 1% NGS, 0.05% Triton X-100 and 0.05% $NaN_3$ in PBS. Samples were washed at least three times for 10-20 minutes each in PBS. The respective anti-mouse and anti-rabbit secondary antibody (Jackson ImmunoResearch, West Grove) conjugates were added in incubation solution for 2 hours at RT. Samples were then finally washed three more times with DNA-PAINT buffer before imaging. Red fluorescent carboxylate-modified microspheres, with diameter of 200 nm (F8887, ThermoFisher Scientific) were added to samples prior to super-resolution imaging to provide fiducials for correcting x-y drift. After imaging D1 sites with P1 imager at a concentration of 0.1-0.5 nM, S1 strands were introduced at a concentration of 500 nM, and P1 binding events monitored as they were suppressed due to S1 binding to D1. After 15 minutes, excess S1 was washed out and replaced with P5 imagers at 0.3 nM to sample D2. Following this, a pre-mixed DNA-PAINT buffer solution containing P1 imager (capable of sampling an open S1-loop), S2 at concentration 500 nM and the blocking strand B at 2-3 µM were introduced, so that S2-B complexes bind D2, making the P5 docking sites unavailable. After 15 minutes, excess unbound S2 were washed out with DNA-PAINT buffer containing P1 imager in a series of washes lasting 15 minutes. Finally, the shield-removal strand was added at ~1 µM and PD-PAINT (S1-loop) signal sampled.

**1.8 Ratio-PD-PAINT imaging of biological samples**

Cardiac cell samples were prepared on coverslips as previously stated. For imaging, we introduced DNA-PAINT buffer containing both P1 and P5 imagers modified to have ATTO 655 and ATTO 700 dyes, respectively. Both dyes were excited simultaneously using a 642 nm CW diode laser and their emission further separated in a splitter device (OptoSplit II, Cairn Research) containing a dichroic mirror (T710 LPXXR-UF3, Chroma). Each of the two images were focused onto separate halves of an iXon Ultra EMCCD (Andor) camera and the two channels aligned as described previously.[8] Once both targets had been sampled S1 was introduced, followed by S2+B and the remover strand. The open loop was finally sampled with P1 ATTO 655.

**1.9 PD-PAINT Imaging of DNA origami samples**

Coverslips were sonicated in acetone for 10 minutes, allowed to air dry and then sonicated in isopropanol for a further 10 minutes before being rinsed with deionized water and air dried. The coverslips were then incubated with a PBS solution of biotin-labeled bovine albumin (A8549, Sigma, concentration 1 mg $mL^{-1}$) for 5 minutes, washed 3 times with PBS and subsequently incubated with a NeutrAvidin (31000, ThermoFisher) solution in PBS, concentration 1 mg $mL^{-1}$, for 5 minutes. Samples were then washed with PBS containing 10 mM magnesium chloride. DNA-origami solutions, with an origami concentration of ~1 nM, were deposited onto the coverslips previously labeled with NeutrAvidin and allowed to attach thanks to the biotin modifications on the origami. After allowing 5 minutes for attachment, samples were again washed to remove any unbound origami. Imaging was carried out using 2 nM P1 imagers in DNA-PAINT buffer. The first phase consisted of detecting and counting accessible origami tiles, targeting the D1 sites using P1, acquiring ~40k frames. S1 at ~500 nM was then added and given at least 15 minutes to fully attach before being washed with DNA-PAINT buffer containing 2 nM P1. Pre-mixed S2 at 500 nM and B at 2 µM was then added. After a further 15 minutes the excess strands were removed through a series of washes before finally introducing R at ~500 nM and acquiring an additional 100k frames. The pre-/post- wash direct imaging control results (Fig. 3b) were obtained using D1 tiles. P1 imager at 2 nM was used for the pre-wash direct phase and was washed out using plain DNA-PAINT buffer. This was replaced with fresh P1 imager of the same concentration ~15 minutes later and allowed to reach an equilibrium for the post-wash direct second imaging phase. Individual control experiments were conducted with origami tiles featuring D1 attachments sites. To these, one or more of S1, S2, S2+B, or R strands were added into solution at concentrations outlined above in order to confirm they functioned as expected, see Figure S6 for detailed event time traces.

**1.10 Imaging setup**

Images were acquired on a modified Nikon Eclipse Ti-E inverted microscope (Nikon, Tokyo) housing a 60× 1.49NA APO oil immersion TIRF objective (Nikon, Tokyo) with an Andor Zyla 4.2 sCMOS (scientific complementary metal-oxide-semiconductor) camera (Andor, Belfast). The ratio-metric data was collected using an iXon Ultra EMCCD camera (Andor) mounted on another port of the same microscope. A piezo objective scanner (P-725, Physik Instrumente, Karlsruhe) was used for focus control. For imager excitation in super-resolution imaging, a 642 nm CW diode laser (Omikron LuxX, Rodgau) was used, at a power of ~15 mW distributed over an illumination spot of approximately 30 µm in diameter. Widefield fluorescence images were excited with a tunable LED-light source (CoolLED, Andover). While thermal drift was reduced with a custom objective holder, any residual focal drift was compensated by continuous feedback from an auxiliary camera in transmission mode at a non-interfering wavelength,[4,9] similar to a method described by McGorty et al.[10] Prior to imaging, 200 nm red fluorescent carboxylate-modified microspheres (F8887, ThermoFisher Scientific) were added at a dilution of ~1:50k in DNA-PAINT buffer, and allowed to settle on the coverslip to serve as fiducials. ROIs were carefully selected to incorporate at least one fiducial in order to correct lateral drift in post-analysis.

**1.11 Image acquisition and analysis**

Synthetic origami samples and isolated myocytes were imaged in Total Internal Reflection Fluorescence (TIRF) mode with a focus just above the coverslip, tissue sections were imaged in Highly Inclined and Laminated Optical sheet (HILO) mode, focusing approximately 1 µm into the tissue. Imager concentrations were 2 nM for origami imaging, and 0.1-0.5 nM for imaging of tissue sections. The entire imaging process, including hardware control, localization and fitting and post-processing was carried out using software in the Python Microscopy Environment (PyME), available freely via: https://bitbucket.org/christian_soeller/python-microscopy-exeter/. Camera frame integration time for super-resolution image acquisition was set to 100 ms. For localization, individual binding events were detected and fitted to a 2D Gaussian model. Out-of-focus binding events were suppressed in a post-processing step by filtering with respect to parameters of the fitting, e.g. localization errors and photon number per event. Binding events which were detected over multiple subsequent frames were merged and the images rendered by jittered triangulation or Gaussian rendering.[11]

**1.12 Assessment of site loss and addressability on origami**

Gaussian rendered super-resolution images from the (pre-wash) direct phase (~40k frames) were used to identify tiles with complete addressability (all 6 sites visible). Images were then rendered utilizing ~100k frames from the PD phase or the post-wash direct phase, and the initially identified complete tiles inspected to count the visible sites and calculate estimates of the fraction of retained sites, $f_R$ and produce the histograms in Fig. 3. Assessment of the of the addressability of D1 and D2 was carried out by counting binding sites from rendered images generated using 38k frames each. Tiles were first visually identified, and then the number of visible sites on each counted to calculate the percentage of addressable sites in Fig. S1. Note that since origami with 1 or 2 addressable sites could hardly be identified by visual inspection against background events, the addressability we estimate with this procedure are likely overestimates. Also note that site D2 could not be directly targeted with P5 imagers on the origami, possibly due to weak non-specific binding of this overhang on the surrounding origami. To overcome this limitation we introduced strand D3 which irreversibly binds to D2, and features a docking site which would be imaged with P1.

For the analysis in Fig. S4, images were rendered from the frame sequence as indicated (e.g. 66K or 33K frames). In the rendered images spots on origami were detected using the "blob finding" functionality in the dh5view application that is part of the PYME software suite, using a signal-to-noise based threshold of 0.4. Detected spots were then matched between images from different frame periods and it was programmatically counted which fraction of spots were retained in imagers from later frame sequences.

**1.13 Kinetic analysis of event time courses of single origami spots**

To investigate the kinetics and thermodynamics of the PD-PAINT platform with single-molecule resolution we extracted the event time courses of individual spots on origami, examples of which are shown in Fig. 4c. First, we corrected any residual drift over the acquisition period using tracked fiducials (see sample preparation above). Following drift correction, non-specific binding locations were identified and discarded as those were binding events are not observed repeatedly over time as expected for proper docking sites.[12,13] The remaining specific events were rendered using Gaussians[11] into separate images for the (pre-wash) direct phases and the PD phases. In the rendered images spots on origami were detected using the "blob finding" functionality in the dh5view application that is part of the PYME software suite, using a signal-to-noise based threshold of 0.4. Detected spot locations were used to generate a label mask that assigns a unique label to a circular area ~60 nm diameter around spot centers with a watershed segmentation procedure implemented in the dh5view application. Using the resulting label mask, all events in a uniquely labeled region were grouped and times of these event groups extracted for further analysis, namely the study of dark-time statistics and qPAINT analysis.

**1.14 qPAINT analysis**

To evaluate the possibility of combining qPAINT analysis with PD-PAINT experiments, we used the time-series of bright events recorded on individual origami sites, from both direct phases and PD phases, to produce the qPAINT calibration curves in Fig. S10. Pseudo-experimental event traces corresponding to $N$=1, 2,…10 emitters were generated by merging $N$ different single-site time series (see Fig. 4). To account for the gradual S1-S2 dimerization transition we restricted single-site traces from the PD phases to the later part of the experiment, after 80% of the detected sites have already produced at least one blink.

Dark periods shorter than 2 seconds (20 frames) were eliminated from the merged time traces as we also do for single-emitter traces, since these very brief dark periods are believed not to correspond to (complete) imager detachment but rather be generated by detection noise and dye blinking. To eliminate short dark times, intervals shorter than 2 seconds are merged into a single event. To avoid oversampling, each single-site time series was used only once for each value of *N*. Therefore, more independent pseudo-experimental multi-site traces could be generated for smaller values of *N*.
For each of the independent pseudo-experimental traces we calculated the mean dark time $\tau_D$. The box plots in Fig. S10 show the distribution of $\tau_D^{-1}$ among the artificially generated ensemble of *N*-site traces. A linear increase of $\tau_D^{-1}$ as a function of *N* was found, as required for the application of qPAINT. Values of the gradient of a least-squares linear fit are given in SI Table S1 with errors from square root of inverse Hessian in optimization. The lower gradient in the PD phases indicates that at any one time not all of the S1 loops are in the open configuration, emitting detectable bright events.

#### 1.15 Analysis of first passage times

After addition of the shield remover, datasets collected in the PD phase of PD-PAINT experiments demonstrate a gradual rise in event rate. We have estimated the timescale of this gradual rise by evaluating, on origami test samples with $d$=5, 10 and 15 nm, the mean time interval between the introduction of the shield-remover strand and the detection of the first bright event on each spot, a quality that we refer to as first-passage-time, $t_{\text{fp}}$. More specifically, $t_{\text{fp}}$ represents the time of first passage from a state where S1 forms a loop, and S2 is bound to a blocking strand, to a state where S1 is bound to S2 and the S1 loop is open, followed by binding to an imager and subsequent detection (see Fig. 1). Cumulative distribution functions of $t_{\text{fp}}$ recorded for $d$=5, 10 and 15 nm are shown in Fig. S3. In all cases, the mean first-passage time $<t_{\text{fp}}>$ is smaller than 1/3 of the duration of the PD phase. Values of $<t_{\text{fp}}>$ are reported in Table S2.

#### 1.16 Single-site kinetic and thermodynamic analysis of the blinking behavior

We analysed the statistics of dark times, time intervals between subsequent blinks detected on a single origami site, to gather information of the kinetics and thermodynamics of the various molecular processes underlying the PD-PAINT platform. As graphically summarised in Fig. 1 and 4, relevant processes include the dimerization and un-binding transition between S1 and S2 probes, described as first-order processes with rate constants $k_d$ and $k_u$, respectively and the second-order process of an imager binding to an exposed docking site, with rate constant $k_{on}$. The off rate of the imager-docking interactions, $k_{off}$ is much larger compared to all other processes (i.e. blinks are very brief compared to the typical dark time).
Single-site event time series are extracted for all origami experiments, both PD-PAINT and the conventional DNA-PAINT control, and for all experimental phases (pre- and post-wash direct phase PD-phases). As discussed above (in the qPAINT section), data was pre-processed by eliminating dark events which were shorter than 20 frames (2 seconds). Cumulative Distribution Functions (CDF) of the dark times were then extracted for visualization of the distribution, as shown in Fig. 4 and Fig. S2.
Two different approaches were then implemented to extract the kinetic rate constants.

*Analytical fit of the dark time distribution*

For conventional DNA-PAINT, as carried out in the pre- and post-wash direct phases of the of the control experiment (Fig. 3a) and in the direct phases of DNA PAINT experiments, the CDF is expected to follow the exponential trend:

$$\mathrm{CDF}(t) = 1 - e^{-k_{on}c\,t}. \qquad \text{Eq. S 4}$$

Here, $c$ is the imager concentration. This model was used to extract $k_{on}c$ values from direct imaging phases through a maximum-likelihood estimation, and the outcomes are listed in Table S3. Uncertainties were calculated via bootstrap resampling of the single molecule traces (100 re-samplings), and quoted the standard deviation of the optimized parameters. Comparison of the experimental CDFs and the respective fits is shown in Fig. S2a.

The blinking behaviour of a PD-PAINT site can instead be modelled as an Interrupted Poisson Process (IPP), which is known to generate independent waiting times according to the hyper-exponential distribution,[14] H2:

$$\mathrm{CDF}(t) = 1 - pe^{-\gamma_1 t} - (1-p)e^{-\gamma_2 t}. \qquad \text{Eq. S 5}$$

Here, the parameters of $p$, $\gamma_1$, and $\gamma_2$, are related to the physical rate constants as

$$p = \frac{1}{2}\frac{k_{on}c - k_u - k_d + \sqrt{(k_{on}c + k_u + k_d)^2 - 4k_{on}ck_d}}{\sqrt{(k_{on}c + k_d + k_u)^2 - 4k_{on}ck_d}}, \qquad \text{Eq. S 6}$$

$$\gamma_1 = \frac{1}{2}\left(k_{on}c + k_d + k_u + \sqrt{(k_{on}c + k_u + k_d)^2 - 4k_{on}ck_d}\right), \qquad \text{Eq. S 7}$$

$$\gamma_2 = \frac{1}{2}\left(k_{on}c + k_d + k_u - \sqrt{(k_{on}c + k_u + k_d)^2 - 4k_{on}ck_d}\right). \qquad \text{Eq. S 8}$$

As above, we have estimated the optimum values for the rate constants via maximum likelihood, and the results are listed in Table S3. The experimental CDF are compared to the fitted analytical model in Fig. S2b, showing excellent agreement.

*Explicit Markov Modelling*

The empirical single molecule traces are only sampled for finite time, which means that empirically observed long dark times will occur less frequently than compared to the H2 distribution used for the analytical fit (Eq. S5). Certainly, no dark event can exceed the duration of the experiment, but also with each subsequent bright event, the time left for the next one to occur is shorter. This means that the analytical fits may be subject to bias which will be dependent on the experimental duration.

To correct for this bias, we have performed explicit Hidden Markov Model simulations of the relevant molecular processes for all PD-PAINT experiments with *d* = 5, 10, and 15 nm.
The analytical fits of $k_{on}c$ are more reliable than those for $k_d$, $k_u$, given that the value of the imager on-rate influences more strongly the short-times portion of the CDF, which is the least affected by finite-sampling artefacts. Therefore, when simulating each PD-PAINT experiment, we have used the $k_{on}c$ value acquired via the analytic fit (Table S3). The rate imager-docking off rate $k_{off}$ is set to a high value, to reflect the fact that bright events are much shorter than the mean dark times, as required in all DNA-PAINT experiments.

With the aim of optimizing for the dimerization and unbinding rates we simulated the system for 400 different $k_d$, $k_u$ pairs, with each parameter taking 20 equally spaced values between $2 \times 10^{-4}\ \text{s}^{-1}$ and $2 \times 10^{-3}\ \text{s}^{-1}$ (inclusive). For each combination of $k_d$ and $k_{u\,,}$ and for each PD-PAINT experiment, we generated 16 sets of Markov traces using the Gillespie algorithm (i.e. 16 Markov trajectories per single molecule trace per parameter combination). To further increase the similarity with the experimental scenario, each Markov simulation was initialized at a time equal to that of the first blink of a randomly selected experimental event trace, and then allowed to run for the reminder of the PD phase.

The simulated event traces are then processed according to the same procedure used for experimental data. Simulated dark times are then pooled between all replicas for a given parameter pair and for a given experiment and binned to create a discrete probability distribution of dark times. The left hand side of bins are uniformly spaced between 0 and 0.97 hours, with 20 intervals, and all bins are considered to be bounded by the left hand side of the bin to the right, except the final one which is treated as half open.

To identify which Markovian transition parameters lead to good approximations of the data, dark times from the experiments are binned in a similar manner to the simulated data. The likelihood of the experimental bins is then calculated given the empirical probability distribution implied by the simulated bins.

The estimates of $k_d$, $k_u$, as summarized in Table S4 are obtained by optimizing the likelihood. Uncertainties on the parameters are evaluated by bootstrap resampling single traces from the experimental datasets, and finding the maximum likelihood from each, with resampling performed 50 times. Uncertainties here are given as the standard deviation of locations of maximum likelihood over resampling events. In two resampling timeseries, the likelihood grid did not have a local maximum, and these samples were not used in the analysis.

Comparison between the experimental dark-time CDFs and simulated equivalents corresponding to the optimized kinetic rates are shown in Fig. S2c.

### 2 Simulation Methods

#### 2.1 Coarse-grained model

We used the coarse-grained model oxDNA to estimate the free energies of S1-S2 hybridization (Fig. 2a, b, main text) and of the imager interacting with the binding site on S1 (Fig. 2c, main text).[15,16] The oxDNA representation has been developed for studying DNA nanostructure reconfiguration in the context of strand displacement reactions[15,17] but is remarkably versatile, accurately modelling the configurational freedom of large origami nanostructures.[18] Each nucleotide is modelled as a bead with three sites, representing the phosphate backbone, the stacking site, and the hydrogen bonding site. Nucleotides interact with each other via anisotropic potentials encoding excluded volume, nucleotide stacking, cross-stacking, and backbone connectivity. Coulomb repulsion between the negatively-charged backbone sites is modelled through the Debye Hückel approximation. The model has been parameterized top down to match a diverse range of thermodynamic and structural features observed in experiments.

#### 2.2 Calculation of S1-S2 dimerization free energy

*Sampling.*

Simulations were run with the oxDNA2 force field, at 25ºC and with a Debye length equivalent to that of a 0.5 M monovalent salt solution, comparable to experimental conditions. Force-field parameters averaged over nucleotides are used. Both geometries shown in Fig. 1 and 2 (main text) were simulated: geometry 1, in which S1 and S2 are directly anchored and the experimentally relevant geometry 2 in which S1 and S2 are anchored via strands D1 and D2. Those nucleotides which in and experiment would be chemically immobilized, namely end points on of S1/S2 in geometry 1 and D1/D2 in geometry 2 are anchored via stiff harmonic springs to points in space separated by distance $d$. We describe those springs as tethers.

For each value of *d*, the free energy difference between the dimerized and un-dimerized states of the S1-S2 system was acquired via umbrella sampling,[19] using multiple thermodynamic windows. Each window was sampled using Virtual Move Monte Carlo (VMMC), a method which is especially suited for the sampling of stiff polymers which otherwise suffer from low acceptance probabilities.[20] The Weighted Histogram Analysis Method (WHAM),[21] as implemented in Python, was used to combine the probability distributions of different thermodynamic windows.

While in reality the opening process of the hairpin on S1 would almost certainly proceed along a branch migration trajectory in line with previously well studied strand displacement reactions,[17] such a trajectory is difficult to sample adequately. Instead, we sampled along an unphysical path where the S1 stem loop opens completely before any of the S1-S2 bonds are formed. More specifically, for each value of $d$, the free energies associated to the formation of the S1 stem-loop and to the formation of the S1S2 dimer were calculated relative to the fully non-bonded state. These were then combined to work out the free energy difference between the S1 stem loop and the S1S2 dimer and the associated dimerization probability, shown in Fig. 2a,b.

The umbrella sampling windows were defined with respect to a on a reaction coordinate, which in turn depends on two observables: the minimum distance $x$ between any two nucleotides which would be hybridized in either the closed S1-loop configuration or when S1 and S2 are hybridised, and the number of formed base pairing bonds $n_B$.. It is convenient to further define binned intervals for $x$ and associate to them a discrete index $m$, as given in Table S10.

Umbrella sampling windows we used are thus defined in Table S11, where the last two columns indicate the use of given windows for calculating the formation free energy of the S1 stem-loop and the S1S2 dimer. Sampling within each of the windows was run initially with biases chosen from experience, and later optimized to ensure flat histogram sampling.

For the geometry 1, tether separation distances of $d$={2, 4, 6, 10, 12, 13, 14, 15} nm were used. The simulation was equilibrated over $10^6$ Monte Carlo steps, and data collected between $1.3 \times 10^6$ and $5.5 \times 10^6$ steps. For geometry 2, with a double stranded extensor, tether separation distances of $d$={5, 10, 15, 20, 25, 30} nm were used. The simulation was run for between $7.91 \times 10^5$ steps and $9.17 \times 10^5$ timesteps depending on the window. One to five repeats were used for each window.

In both cases, the state of the system (reaction coordinate) was sampled every 1000 steps. For evaluation of errors, the reaction-coordinate timeseries for each window were concatenated over repeats and then divided equally into three parts. WHAM was performed on the sets individually (see below). Errors are then stated as the standard error over those sets.

For evaluating the formation free energy of the S1 stem-loop we assume that the latter is formed when at least one S1-S2 base pairing bond is present. Likewise, for the case of the S1-S2 dimerization free energy, we assume that a dimer is formed whenever S1-S2 share a bond.

*WHAM*

The Weighted Histogram Analysis Method (WHAM) is implemented to collate probability distributions sampled within different windows, and relies on solving the coupled equations

$$p_j = \frac{\sum_{i=1}^{S} n_{ij}}{\sum_{i=1}^{S} N_i f_i c_{ij}}, \qquad \text{Eq. S 9}$$

$$f_i^{-1} = \sum_{j=1}^{M} c_{ij}\, p_j. \qquad \text{Eq. S 10}$$

Here, the suffix $i$ indexes over the $S$ windows, and $j$ indexes over the $M$ configurations (values of the order parameter). We seek to extract $p_j$, which are the probabilities of finding the system in configuration $j$, while $f_i$ are normalizing factors which may take a different value in each window. $n_{ij}$ is the number of samples of configuration j in window $i$, as sampled from MC runs, while $N_i$ is the total number of samples from window $i$, and $c_{ij}$ is the bias of configuration $j$ in window $i$.
To extract $p_j$ we solve Eq. S 9 and Eq. S 10 iteratively: first we use the initial guess that $p_j$ are uniform to generate $f_i$ from Eq. S 10, then using the values of $f_i$ to make a better approximation to $p_j$ with Eq. S9 , and so forth until convergence.

**2.3 S1-imager dimerization free energy**

To extract the hybridization free energy of the imager P1 to docking sites, shown in Fig. 2c, we use sequence-dependent interaction potentials. In contrast to simulations of S1-S2 hybridization, misbonding between nucleotides is permitted. i.e. any A can bind to any T and any C to any G regardless of whether they are expected to do so in the target configuration.
For these calculations we use geometry 1 (Fig. 1 and 2), namely S1 and S2 are directly anchored via a polyT spacer. Additionally, different from the experimental implementation, here the number of base-pairs in the hairpin loop of S1 is 20 rather than 14. This difference is likely to have no impact, given that it occurs far from the binding site and that a 14 base-pair loop is highly unlikely to open over under reaction conditions, similarly to the 20 base-pair loop.

Simulations were run to evaluate the free energy $\Delta G$ of imager P1 binding to the docking domain on S1 in three cases, as discussed in the main text (Fig. 2b):

i) Closed S1 loop, where the stem of the hairpin on S1 is allowed to close. Here only S1 is present, with sequence given in Table S6 as S1''.
ii) Open S1 loop, where the S1-S2 dimer is forced to form using a potential which requires that at least 17 of the 20 S1-S2 bonds are formed. The sequences used for S1 and S2 are given in Table S6 as S1'', and S2'.
iii) Conventional DNA-PAINT configuration, using a different S1-sequence in which loop formation is prevented (see sequence Classical PAINT S1' in Table S6)

We follow a simulation protocol analogous to what was done for the S1-S2 interaction free energy. Specifically, we perform umbrella sampling using a set of windows defined in terms of a two-dimensional reaction coordinate $\vec{q} = (b, d)$, where $b$ is the number of P1-S1 bonds and $d$ is a discrete measure of distance between P1 and its complementary domain on S1, defined as explained above for the S1-S2 case. The chosen windows are here listed in Table S11. Sampling within each window is performed using the VMMC algorithm[20] and the probability distributions from each window stitched together using WHAM.[21] We simulated our system in a periodic cubic box with size of 85 nm, corresponding to an effective concentration of $c = 2.7$ µM for each of the strands.
By defining as $p(b)$ the probability of P1 forming $b$ bonds with S1, we extract

$$\Delta G_{\text{S1-P1}}(b) = \begin{cases} -k_\text{B}T \log\left[\dfrac{p(b)}{1-p(b)}\right] \text{for b=0} \\ -k_\text{B}T \log\left[\dfrac{p(b)}{1-p(b)}\dfrac{c_\text{exp}}{c}\right] \text{for b>0,} \end{cases}$$ Eq. S 11

where the correction applied for $b > 0$ accounts for the different imager concentration $c_\text{exp}$ used in experiments, where $c_\text{exp} = 2$ nM.

To extract the timescale for the dissociation of the imager we evaluated the probability of P1 and S1 having at least one base-pairing bond, $p_\text{bound} = \sum_{b=1}^{9} p(b)$. Then, the equilibrium constant was computed as

$$K = \frac{p_\text{bound}}{1-p_\text{bound}}\frac{\rho_0}{c}$$ Eq. S 12

$\rho_0$ is a reference concentration, here, 1M. The P1-S1 off rate can then be estimated as $k_\text{off} = k_\text{on}/K$. Here we used the approximate estimate $k_\text{on} = 10^6\text{M}^{-1}\text{s}^{-1}$, a value taken from experimental studies,[22] and thought to be accurate for sufficiently short oligonucleotides at sufficiently high ionic strengths, conditions that should be fulfilled in our experimental system. The P1-S1 binding lifetimes are then extracted as $\tau_\text{bound} = 1/k_\text{off}$.

**Supplementary Tables**

**Table S1:** qPAINT inverse mean dark times $\tau_D^{-1}$ per emitter, as fitted from pseudo-experimental calibration curves shown in Fig. S10. Uncertainties are given as the square root of diagonal entries in the inverse Hessian of a least squares fit.

| $d$ | Direct phase $\tau_D^{-1} \times 10^{-3}\ (s^{-1})$ | PD phase $\tau_D^{-1} \times 10^{-3}\ (s^{-1})$ |
|---|---|---|
| 5 nm | 3.80 ± 0.02 | 2.11 ± 0.01 |
| 10 nm | 2.96 ± 0.01 | 1.76 ± 0.02 |
| 15 nm | 3.10 ± 0.02 | 1.61 ± 0.01 |

**Table S2:** Mean first passage times for bright events to be regenerated in PD-PAINT for the different DNA S1-S2 distances. A first passage times are defined as the time interval between the addition of the remover DNA strand, and the first bright event, and the mean first passage time is calculated from the mean across different spatially resolved single molecule timeseries. See Fig. S3.

| $d$ | FPT (minutes) |
|---|---|
| 5 nm | 28 |
| 10 nm | 48 |
| 15 nm | 45 |

**Table S3**: Estimates of the rates of the process of imager hybridization as determined from exponential fits of the dark time distributions in direct phases, namely by targeting D1 with imager P1 as show in Fig. 3a. The experimental and theoretical CDFs are shown in Fig. S2a. Here $k_{on}$ is the second-order rate constant and $c$ = [P1] = 2 nM. All uncertainties are calculated through bootstrap resampling.

| Origami System | $k_{on}c \times 10^{-3}\ (s^{-1})$ |
|---|---|
| PD-PAINT 5 nm | 5.01 ± 0.14 |
| PD-PAINT 10 nm | 3.76 ± 0.12 |
| PD-PAINT 15 nm | 4.07 ± 0.10 |
| Control (pre) | 2.94 ± 0.05 |
| Control (post) | 3.10 ± 0.06 |

**Table S4:** Analytical estimates of the rates of the processes underlying the blinking kinetics of PD-PAINT sites, where $k_d$ and $k_u$ are the first-order rate constants of S1-S2 dimerization and unbinding, $k_{on}$ is the second-order rate constant of the imager binding to the S1 socking site, and $c$ the imager concentration. The P1-S1 reaction rate $k_{on}c$ is fitted as a single parameter. Estimates for these parameters are evaluated from a maximum likelihood fit of the waiting times for an Interrupted Poisson Process to the observed dataset of waiting time events. See SI text and Fig. S2a. From the estimated $k_d$ and $k_u$, we can extract the equilibrium constant of the S1-S2 dimerization reaction, the dimerization probability and the corresponding free energy ΔG. All uncertainties are calculated through bootstrap resampling.

| $d$ | $k_{on}c \times 10^{-3}\ (s^{-1})$ | $k_d \times 10^{-3}(s^{-1})$ | $k_u \times 10^{-3}(s^{-1})$ | K | Prob S1-S2 dimer | ΔG ($k_BT$) |
|---|---|---|---|---|---|---|
| 5 nm | 5.5 ± 0.2 | 1.10 ± 0.11 | 0.70 ± 0.10 | 1.6 ± 0.2 | 0.61 ± 0.02 | -0.47 ± 0.10 |
| 10 nm | 4.90 ± 0.2 | 1.40 ± 0.19 | 0.80 ± 0.18 | 1.8 ± 0.3 | 0.65 ±0 .03 | -0.64 ± 0.14 |
| 15 nm | 4.94 ± 0.2 | 1.18 ± 0.12 | 0.89 ± 0.15 | 1.36 ± 0.15 | 0.58 ± 0.03 | -0.31 ± 0.12 |

**Table S5:** Numerical estimates of the rate constants described in Table S3 as extracted via explicitly replicating the experiment using a Hidden Markov Model. See SI Text and Fig. S2b.

| $d$ | $k_d \times 10^{-3}(s^{-1})$ | $k_u$- $\times 10^{-3}(s^{-1})$ | K | Prob S1-S2 dimer | ΔG ($k_BT$) |
|---|---|---|---|---|---|
| 5 nm | 1.21 ± 0.2 | 0.80 ± 0.14 | 1.5 ± 0.3 | 0.60 ± 0.04 | -0.41 ± 0.17 |
| 10 nm | 1.4 ± 0.2 | 0.95 ± 0.15 | 1.5 ± 0.3 | 0.59 ± 0.04 | -0.38 ± 0.18 |
| 15 nm | 1.33 ± 0.16 | 1.2 ± 0.2 | 1.16 ± 0.17 | 0.53 ± 0.03 | -0.14 ± 0.14 |

**Table S6:** PD-PAINT DNA sequences used in DNA origami, biological experiments and molecular simulations (5' → 3').

| **Name** | **Sequence** |
|---|---|
| D1 | [AzideN] TTA TAC ATC TAT TTC TTC ATT ATT CAC TTA CTA [Cy3] |
| D2 | [AzideN] TTT TAG GTA AAT TTT GAT TGT GAG GAA G [Cy5] |
| P1 imager | CTA GAT GTA T [At655] |
| P5 imager | CTT TAC CTA A [At655] or CTT TAC CTA A [At700] |
| S1 | AGG AGA GGA GAA TAC ATC TAT ATT CTC CTC TCC TCC TTC CTT TTT TTT TTT TTT TTA GTA AGT GAA TAA TGA AGA AAT AGA TGT ATA A |
| S2 | CTT CCT CAC AAT CAA AAT TTA CCT AAA ATT TTT TTT TTT TTT TGG AAG GAG GAG AGG AGA ATA |
| S2 shield | TCT TCA TTA CCG AGC GTA TCC TCC TTC CAA AAT TGT CTT GTA TGA T |
| S2 shield remover | ATC ATA CAA GAC AAT TTT TTT TTT TTG ATA CGC TCG GTA ATG AAG A |
| S1' | AGG AGA GGA GAA TAC ATC TAT ATT CTC CTC TCC TCC TTC CTT TTT TTT TTT TTT TT [Harmonic Restraint] |
| S2' | [Harmonic Restraint] TT TTT TTT TTT TTT TGG AAG GAG GAG AGG AGA ATA |
| S1'' | GGA AGG AGG AGA GGA GAA TAC ATC TAT ATT CTC CTC TCC TCC TTC CTT TTT TTT TTT TTT T [Harmonic Restraint] |
| Classical Paint S1 | GGA AGG AGG AGA GGA GAA TAC ATC TAT ATT GTG GTG TGG TGG TTG GTT TTT TTT TTT TTT T [Harmonic Restraint] |
| D3 | ATA CAT CTA ATA CAT CTA ATA CAT CTA ATA CAT CTA ATA CAT CTA CTT CCT CAC AAT CAA AAT TTA CCT AAC ATA CAT CTA ATA CAT CTA ATA CAT CTA ATA CAT CTA ATA CAT CTA |

**Table S7:** DNA sequences for DNA origami synthesis – sequences with 5' biotin end modifications (5' → 3').

| **Oligo name** | **Sequence** |
|---|---|
| Biotin 1 | /5Biosg/ATTAAGTTTACCGAGCTCGAATTCGGGAAACCTGTCGTGC |
| Biotin 2 | /5Biosg/ATAAGGGAACCGGATATTCATTACGTCAGGACGTTGGGAA |
| Biotin 3 | /5Biosg/GCGATCGGCAATTCCACACAACAGGTGCCTAATGAGTG |
| Biotin 4 | /5Biosg/TTGTGTCGTGACGAGAAACACCAAATTTCAACTTTAAT |
| Biotin 5 | /5Biosg/ATTCATTTTTGTTTGGATTATACTAAGAAACCACCAGAAG |
| Biotin 6 | /5Biosg/CACCCTCAGAAACCATCGATAGCATTGAGCCATTTGGGAA |
| Biotin 7 | /5Biosg/AACAATAACGTAAAACAGAAATAAAAATCCTTTGCCCGAA |
| Biotin 8 | /5Biosg/AGCCACCACTGTAGCGCGTTTTCAAGGGAGGGAAGGTAAA |

**Table S8:** DNA sequences for DNA origami synthesis – sequences without biotin modifications (5' → 3'). Colors of overhang sequences indicate the overhang set in question, either D1 (red), D2a (blue), D2b (green), or D2c (purple). Where a set of overhangs was not used, the strand with the overhang was substituted by a corresponding DNA strand which lacked the colored overhang. Origami were made featuring D1 only (D1 set only), with D1 and D2a (5 nm spacing), with D1 and D2b (10 nm spacing), or with D1 and D2c (15 nm spacing).

| **Oligo name** | **Sequence** | **Overhang set** |
|---|---|---|
| 21[32]23[31]D1 | TTTTCACTCAAAGGGCGAAAAACCATCACC **TTA TAC ATC TAT TTC TTC ATT ATT CAC TTA CTA** | **D1** |
| 19[32]21[31]BLK | GTCGACTTCGGCCAACGCGCGGGGTTTTTC **TTT TAG GTA AAT T TTG ATT GTG AGG AAG** | **D2a** |
| 17[32]19[31]BLK | TGCATCTTTCCCAGTCACGACGGCCTGCAG **TTT TAG GTA AAT T TTG ATT GTG AGG AAG** | **D2b** |
| 15[32]17[31]BLK | TAATCAGCGGATTGACCGTAATCGTAACCG **TTT TAG GTA AAT T TTG ATT GTG AGG AAG** | **D2c** |
| 13[32]15[31]BLK | AACGCAAAATCGATGAACGGTACCGGTTGA | |
| 11[32]13[31]BLK | AACAGTTTTGTACCAAAAACATTTTATTTC | |
| 9[32]11[31]BLK | TTTACCCCAACATGTTTTAAATTTCCATAT | |

| | | |
|---|---|---|
| 7[32]9[31]BLK | TTTAGGACAAATGCTTTAAACAATCAGGTC | |
| 5[32]7[31]BLK | CATCAAGTAAAACGAACTAACGAGTTGAGA **TTT TAG GTA AAT T TTG ATT GTG AGG AAG** | **D2c** |
| 3[32]5[31]BLK | AATACGTTTGAAAGAGGACAGACTGACCTT **TTT TAG GTA AAT T TTG ATT GTG AGG AAG** | **D2b** |
| 1[32]3[31]BLK | AGGCTCCAGAGGCTTTGAGGACACGGGTAA **TTT TAG GTA AAT T TTG ATT GTG AGG AAG** | **D2a** |
| 0[47]1[31]D1 | AGAAAGGAACAACTAAAGGAATTCAAAAAAA **TTA TAC ATC TAT TTC TTC ATT ATT CAC TTA CTA** | **D1** |
| 23[32]22[48]BLK | CAAATCAAGTTTTTTGGGGTCGAAACGTGGA | |
| 22[47]20[48]BLK | CTCCAACGCAGTGAGACGGGCAACCAGCTGCA | |
| 20[47]18[48]BLK | TTAATGAACTAGAGGATCCCCGGGGGGTAACG | |
| 18[47]16[48]BLK | CCAGGGTTGCCAGTTTGAGGGGACCCGTGGGA | |
| 16[47]14[48]BLK | ACAAACGGAAAAGCCCCAAAAACACTGGAGCA | |
| 14[47]12[48]BLK | AACAAGAGGGATAAAAATTTTTAGCATAAAGC | |
| 12[47]10[48]BLK | TAAATCGGGATTCCCAATTCTGCGATATAATG | |
| 10[47]8[48]BLK | CTGTAGCTTGACTATTATAGTCAGTTCATTGA | |
| 8[47]6[48]BLK | ATCCCCCTATACCACATTCAACTAGAAAAATC | |
| 6[47]4[48]BLK | TACGTTAAAGTAATCTTGACAAGAACCGAACT | |
| 4[47]2[48]BLK | GACCAACTAATGCCACTACGAAGGGGGTAGCA | |
| 2[47]0[48]BLK | ACGGCTACAAAAGGAGCCTTTAATGTGAGAAT | |
| 21[56]23[63]BLK | AGCTGATTGCCCTTCAGAGTCCACTATTAAAGGGTGCCGT | |
| 15[64]18[64]BLK | GTATAAGCCAACCCGTCGGATTCTGACGACAGTATCGGCCGCAAGGCG | |
| 13[64]15[63]BLK | TATATTTTGTCATTGCCTGAGAGTGGAAGATT | |
| 11[64]13[63]BLK | GATTTAGTCAATAAAGCCTCAGAGAACCCTCA | |
| 9[64]11[63]BLK | CGGATTGCAGAGCTTAATTGCTGAAACGAGTA | |
| 7[56]9[63]BLK | ATGCAGATACATAACGGGAATCGTCATAAATAAAGCAAAG | |
| 1[64]4[64]BLK | TTTATCAGGACAGCATCGGAACGACACCAACCTAAAACGAGGTCAATC | |
| 0[79]1[63]BLK | ACAACTTTCAACAGTTTCAGCGGATGTATCGG | |
| 23[64]22[80]BLK | AAAGCACTAAATCGGAACCCTAATCCAGTT | |
| 22[79]20[80]BLK | TGGAACAACCGCCTGGCCCTGAGGCCCGCT | |
| 20[79]18[80]BLK | TTCCAGTCGTAATCATGGTCATAAAAGGGG | |
| 18[79]16[80]BLK | GATGTGCTTCAGGAAGATCGCACAATGTGA | |
| 16[79]14[80]BLK | GCGAGTAAAAATATTTAAATTGTTACAAAG | |
| 14[79]12[80]BLK | GCTATCAGAAATGCAATGCCTGAATTAGCA | |

| | |
|---|---|
| 12[79]10[80]BL K | AAATTAAGTTGACCATTAGATACTTTTGCG |
| 10[79]8[80]BLK | GATGGCTTATCAAAAAGATTAAGAGCGTCC |
| 8[79]6[80]BLK | AATACTGCCCAAAAGGAATTACGTGGCTCA |
| 6[79]4[80]BLK | TTATACCACCAAATCAACGTAACGAACGAG |
| 4[79]2[80]BLK | GCGCAGACAAGAGGCAAAAGAATCCCTCAG |
| 2[79]0[80]BLK | CAGCGAAACTTGCTTTCGAGGTGTTGCTAA |
| 21[96]23[95]BL K | AGCAAGCGTAGGGTTGAGTGTTGTAGGGAGCC |
| 19[96]21[95]BL K | CTGTGTGATTGCGTTGCGCTCACTAGAGTTGC |
| 17[96]19[95]BL K | GCTTTCCGATTACGCCAGCTGGCGGCTGTTTC |
| 15[96]17[95]BL K | ATATTTTGGCTTTCATCAACATTATCCAGCCA |
| 13[96]15[95]BL K | TAGGTAAACTATTTTTGAGAGATCAAACGTTA |
| 11[96]13[95]BL K | AATGGTCAACAGGCAAGGCAAAGAGTAATGTG |
| 9[96]11[95]BLK | CGAAAGACTTTGATAAGAGGTCATATTTCGCA |
| 7[96]9[95]BLK | TAAGAGCAAATGTTTAGACTGGATAGGAAGCC |
| 5[96]7[95]BLK | TCATTCAGATGCGATTTTAAGAACAGGCATAG |
| 3[96]5[95]BLK | ACACTCATCCATGTTACTTAGCCGAAAGCTGC |
| 1[96]3[95]BLK | AAACAGCTTTTTGCGGGATCGTCAACACTAAA |
| 0[111]1[95]BLK | TAAATGAATTTTCTGTATGGGATTAATTTCTT |
| 23[96]22[112]B LK | CCCGATTTAGAGCTTGACGGGGAAAAAGAATA |
| 22[111]20[112]B LK | GCCCGAGAGTCCACGCTGGTTTGCAGCTAACT |
| 20[111]18[112]B LK | CACATTAAAATTGTTATCCGCTCATGCGGGCC |
| 18[111]16[112]B LK | TCTTCGCTGCACCGCTTCTGGTGCGGCCTTCC |
| 16[111]14[112]B LK | TGTAGCCATTAAAATTCGCATTAAATGCCGGA |
| 14[111]12[112]B LK | GAGGGTAGGATTCAAAAGGGTGAGACATCCAA |
| 12[111]10[112]B LK | TAAATCATATAACCTGTTTAGCTAACCTTTAA |
| 10[111]8[112]B LK | TTGCTCCTTTCAAATATCGCGTTTGAGGGGGT |
| 8[111]6[112]BL K | AATAGTAAACACTATCATAACCCTCATTGTGA |
| 6[111]4[112]BL K | ATTACCTTTGAATAAGGCTTGCCCAAATCCGC |
| 4[111]2[112]BL K | GACCTGCTCTTTGACCCCCAGCGAGGGAGTTA |
| 2[111]0[112]BL K | AAGGCCGCTGATACCGATAGTTGCGACGTTAG |
| 21[120]23[127]B LK | CCCAGCAGGCGAAAAATCCCTTATAAATCAAGCCGGCG |
| 15[128]18[128]B LK | TAAATCAAAATAATTCGCGTCTCGGAAACCAGGCAAAGGG AAGG |
| 13[128]15[127]B LK | GAGACAGCTAGCTGATAAATTAATTTTTGT |

| | | |
|---|---|---|
| 11[128]13[127]BLK | TTTGGGGATAGTAGTAGCATTAAAAGGCCG | |
| 9[128]11[127]BLK | GCTTCAATCAGGATTAGAGAGTTATTTTCA | |
| 7[120]9[127]BLK | CGTTTACCAGACGACAAAGAAGTTTTGCCATAATTCGA | |
| 1[128]4[128]BLK | TGACAACTCGCTGAGGCTTGCATTATACCAAGCGCGATGATAAA | |
| 0[143]1[127]BLK | TCTAAAGTTTTGTCGTCTTTCCAGCCGACAA | |
| 21[160]22[144]BLK | TCAATATCGAACCTCAAATATCAATTCCGAAA | |
| 19[160]20[144]BLK | GCAATTCACATATTCCTGATTATCAAAGTGTA | |
| 17[160]18[144]BLK | AGAAAACAAAGAAGATGATGAAACAGGCTGCG | |
| 15[160]16[144]BLK | ATCGCAAGTATGTAAATGCTGATGATAGGAAC | |
| 13[160]14[144]BLK | GTAATAAGTTAGGCAGAGGCATTTATGATATT | |
| 11[160]12[144]BLK | CCAATAGCTCATCGTAGGAATCATGGCATCAA | |
| 9[160]10[144]BLK | AGAGAGAAAAAAATGAAAATAGCAAGCAAACT | |
| 7[160]8[144]BLK | TTATTACGAAGAACTGGCATGATTGCGAGAGG | |
| 5[160]6[144]BLK | GCAAGGCCTCACCAGTAGCACCATGGGCTTGA | |
| 3[160]4[144]BLK | TTGACAGGCCACCACCAGAGCCGCGATTTGTA | |
| 1[160]2[144]BLK | TTAGGATTGGCTGAGACTCCTCAATAACCGAT | |
| 0[175]0[144]BLK | TCCACAGACAGCCCTCATAGTTAGCGTAACGA | |
| 23[128]23[159]D1 | AACGTGGCGAGAAAGGAAGGGAAACCAGTAA **TTA TAC ATC TAT TTC TTC ATT ATT CAC TTA CTA** | **D1** |
| 22[143]21[159]BLK | TCGGCAAATCCTGTTTGATGGTGGACCCTCAA **TTT TAG GTA AAT T TTG ATT GTG AGG AAG** | **D2a** |
| 20[143]19[159]BLK | AAGCCTGGTACGAGCCGGAAGCATAGATGATG **TTT TAG GTA AAT T TTG ATT GTG AGG AAG** | **D2b** |
| 18[143]17[159]BLK | CAACTGTTGCGCCATTCGCCATTCAAACATCA **TTT TAG GTA AAT T TTG ATT GTG AGG AAG** | **D2c** |
| 16[143]15[159]BLK | GCCATCAAGCTCATTTTTTAACCACAAATCCA | |
| 14[143]13[159]BLK | CAACCGTTTCAAATCACCATCAATTCGAGCCA | |
| 12[143]11[159]BLK | TTCTACTACGCGAGCTGAAAAGGTTACCGCGC | |
| 10[143]9[159]BLK | CCAACAGGAGCGAACCAGACCGGAGCCTTTAC | |
| 8[143]7[159]BLK | CTTTTGCAGATAAAAACCAAAATAAAGACTCC **TTT TAG GTA AAT T TTG ATT GTG AGG AAG** | **D2c** |
| 6[143]5[159]BLK | GATGGTTTGAACGAGTAGTAAATTTACCATTA **TTT TAG GTA AAT T TTG ATT GTG AGG AAG** | **D2b** |
| 4[143]3[159]BLK | TCATCGCCAACAAAGTACAACGGACGCCAGCA **TTT TAG GTA AAT T TTG ATT GTG AGG AAG** | **D2a** |

| | | |
|---|---|---|
| 2[143]1[159]D1 | ATATTCGGAACCATCGCCCACGCAGAGAAGGA **TTA TAC ATC TAT TTC TTC ATT ATT CAC TTA CTA** | **D1** |
| 23[160]22[176]BLK | TAAAAGGGACATTCTGGCCAACAAAGCATC | |
| 22[175]20[176]BLK | ACCTTGCTTGGTCAGTTGGCAAAGAGCGGA | |
| 20[175]18[176]BLK | ATTATCATTCAATATAATCCTGACAATTAC | |
| 18[175]16[176]BLK | CTGAGCAAAAATTAATTACATTTTGGGTTA | |
| 16[175]14[176]BLK | TATAACTAACAAAGAACGCGAGAACGCCAA | |
| 14[175]12[176]BLK | CATGTAATAGAATATAAAGTACCAAGCCGT | |
| 12[175]10[176]BLK | TTTTATTTAAGCAAATCAGATATTTTTTGT | |
| 10[175]8[176]BLK | TTAACGTCTAACATAAAAACAGGTAACGGA | |
| 8[175]6[176]BLK | ATACCCAACAGTATGTTAGCAAATTAGAGC | |
| 6[175]4[176]BLK | CAGCAAAAGGAAACGTCACCAATGAGCCGC | |
| 4[175]2[176]BLK | CACCAGAAAGGTTGAGGCAGGTCATGAAAG | |
| 2[175]0[176]BLK | TATTAAGAAGCGGGGTTTTGCTCGTAGCAT | |
| 21[184]23[191]BLK | TCAACAGTTGAAAGGAGCAAATGAAAAATCTAGAGATAGA | |
| 15[192]18[192]BLK | TCAAATATAACCTCCGGCTTAGGTAACAATTTCATTTGAAGGCGAATT | |
| 13[192]15[191]BLK | GTAAAGTAATCGCCATATTTAACAAAACTTTT | |
| 11[192]13[191]BLK | TATCCGGTCTCATCGAGAACAAGCGACAAAAG | |
| 9[192]11[191]BLK | TTAGACGGCCAAATAAGAAACGATAGAAGGCT | |
| 7[184]9[191]BLK | CGTAGAAAATACATACCGAGGAAACGCAATAAGAAGCGCA | |
| 1[192]4[192]BLK | GCGGATAACCTATTATTCTGAAACAGACGATTGGCCTTGAAGAGCCAC | |
| 0[207]1[191]BLK | TCACCAGTACAAACTACAACGCCTAGTACCAG | |
| 23[192]22[208]BLK | ACCCTTCTGACCTGAAAGCGTAAGACGCTGAG | |
| 22[207]20[208]BLK | AGCCAGCAATTGAGGAAGGTTATCATCATTTT | |
| 20[207]18[208]BLK | GCGGAACATCTGAATAATGGAAGGTACAAAAT | |
| 18[207]16[208]BLK | CGCGCAGATTACCTTTTTTAATGGGAGAGACT | |
| 16[207]14[208]BLK | ACCTTTTTATTTTAGTTAATTTCATAGGGCTT | |
| 14[207]12[208]BLK | AATTGAGAATTCTGTCCAGACGACTAAACCAA | |

| | |
|---|---|
| 12[207]10[208]BLK | GTACCGCAATTCTAAGAACGCGAGTATTATTT |
| 10[207]8[208]BLK | ATCCCAATGAGAATTAACTGAACAGTTACCAG |
| 8[207]6[208]BLK | AAGGAAACATAAAGGTGGCAACATTATCACCG |
| 6[207]4[208]BLK | TCACCGACGCACCGTAATCAGTAGCAGAACCG |
| 4[207]2[208]BLK | CCACCCTCTATTCACAAACAAATACCTGCCTA |
| 2[207]0[208]BLK | TTTCGGAAGTGCCGTCGAGAGGGTGAGTTTCG |
| 21[224]23[223]BLK | CTTTAGGGCCTGCAACAGTGCCAATACGTG |
| 19[224]21[223]BLK | CTACCATAGTTTGAGTAACATTTAAAATAT |
| 17[224]19[223]BLK | CATAAATCTTTGAATACCAAGTGTTAGAAC |
| 15[224]17[223]BLK | CCTAAATCAAAATCATAGGTCTAAACAGTA |
| 13[224]15[223]BLK | ACAACATGCCAACGCTCAACAGTCTTCTGA |
| 11[224]13[223]BLK | GCGAACCTCCAAGAACGGGTATGACAATAA |
| 9[224]11[223]BLK | AAAGTCACAAAATAAACAGCCAGCGTTTTA |
| 7[224]9[223]BLK | AACGCAAAGATAGCCGAACAAACCCTGAAC |
| 5[224]7[223]BLK | TCAAGTTTCATTAAAGGTGAATATAAAAGA |
| 3[224]5[223]BLK | TTAAAGCCAGAGCCGCCACCCTCGACAGAA |
| 1[224]3[223]BLK | GTATAGCAAACAGTTAATGCCCAATCCTCA |
| 0[239]1[223]BLK | AGGAACCCATGTACCGTAACACTTGATATAA |
| 23[224]22[240]BLK | GCACAGACAATATTTTTGAATGGGGTCAGTA |
| 22[239]20[240]BLK | TTAACACCAGCACTAACAACTAATCGTTATTA |
| 20[239]18[240]BLK | ATTTTAAAATCAAAATTATTTGCACGGATTCG |
| 18[239]16[240]BLK | CCTGATTGCAATATATGTGAGTGATCAATAGT |
| 16[239]14[240]BLK | GAATTTATTTAATGGTTTGAAATATTCTTACC |
| 14[239]12[240]BLK | AGTATAAAGTTCAGCTAATGCAGATGTCTTTC |
| 12[239]10[240]BLK | CTTATCATTCCCGACTTGCGGGAGCCTAATTT |
| 10[239]8[240]BLK | GCCAGTTAGAGGGTAATTGAGCGCTTTAAGAA |
| 8[239]6[240]BLK | AAGTAAGCAGACACCACGGAATAATATTGACG |
| 6[239]4[240]BLK | GAAATTATTGCCTTTAGCGTCAGACCGGAACC |
| 4[239]2[240]BLK | GCCTCCCTCAGAATGGAAAGCGCAGTAACAGT |

| | | |
|---|---|---|
| 2[239]0[240]BLK | GCCCGTATCCGGAATAGGTGTATCAGCCCAAT | |
| 21[248]23[255]BLK | AGATTAGAGCCGTCAAAAAACAGAGGTGAGGCCTATTAGT | |
| 15[256]18[256]BLK | GTGATAAAAAGACGCTGAGAAGAGATAACCTTGCTTCTGTTCGGGAGA | |
| 13[256]15[255]BLK | GTTTATCAATATGCGTTATACAAACCGACCGT | |
| 11[256]13[255]BLK | GCCTTAAACCAATCAATAATCGGCACGCGCCT | |
| 9[256]11[255]BLK | GAGAGATAGAGCGTCTTTCCAGAGGTTTTGAA | |
| 7[248]9[255]BLK | GTTTATTTTGTCACAATCTTACCGAAGCCCTTTAATATCA | |
| 1[256]4[256]BLK | CAGGAGGTGGGGTCAGTGCCTTGAGTCTCTGAATTTACCGGGAACCAG | |
| 0[271]1[255]BLK | CCACCCTCATTTTCAGGGATAGCAACCGTACT | |
| 23[256]22[272]D1 | CTTTAATGCGCGAACTGATAGCCCCACCAG **TTA TAC ATC TAT TTC TTC ATT ATT CAC TTA CTA** | **D1** |
| 22[271]20[272]BLK | CAGAAGATTAGATAATACATTTGTCGACAA **TTT TAG GTA AAT T TTG ATT GTG AGG AAG** | **D2a** |
| 20[271]18[272]BLK | CTCGTATTAGAAATTGCGTAGATACAGTAC **TTT TAG GTA AAT T TTG ATT GTG AGG AAG** | **D2b** |
| 18[271]16[272]BLK | CTTTTACAAAATCGTCGCTATTAGCGATAG **TTT TAG GTA AAT T TTG ATT GTG AGG AAG** | **D2c** |
| 16[271]14[272]BLK | CTTAGATTTAAGGCGTTAAATAAAGCCTGT | |
| 14[271]12[272]BLK | TTAGTATCACAATAGATAAGTCCACGAGCA | |
| 12[271]10[272]BLK | TGTAGAAATCAAGATTAGTTGCTCTTACCA | |
| 10[271]8[272]BLK | ACGCTAACACCCACAAGAATTGAAAATAGC | |
| 8[271]6[272]BLK | AATAGCTATCAATAGAAAATTCAACATTCA **TTT TAG GTA AAT T TTG ATT GTG AGG AAG** | **D2c** |
| 6[271]4[272]BLK | ACCGATTGTCGGCATTTTCGGTCATAATCA **TTT TAG GTA AAT T TTG ATT GTG AGG AAG** | **D2b** |
| 4[271]2[272]BLK | AAATCACCTTCCAGTAAGCGTCAGTAATAA **TTT TAG GTA AAT T TTG ATT GTG AGG AAG** | **D2a** |
| 2[271]0[272]D1 | GTTTTAACTTAGTACCGCCACCCAGAGCCA **TTA TAC ATC TAT TTC TTC ATT ATT CAC TTA CTA** | **D1** |

**Table S9:** Expected binding timescales for the dissociation of the imager (P1), to the docking region on S1. Uncertainties shown here were calculated considering only the uncertainty in the hybridization equilibrium constant *K* evaluated from simulation. (*) This estimate is an approximate lower bound, as here, the on-rate $k_{on}$ used in the calculation of the timescale from the equilibrium constant *K* (obtained from the simulations) is that of simple hybridizing strands ($10^6$ $M^{-1}$ $s^{-1}$), whereas the loop will limit the rate of encounter between the binding sites.

| System | Estimated Timescale of Binding |
|---|---|
| S1 closed loop | 2.0 ± 0.4 μs * |
| S1 open loop | 0.27 ± 0.07 s |
| P1 docking (classical DNA-PAINT) | 0.19 ± 0.03 s |

**Table S10:** Correspondence of distances in free energy of opening simulations to binned order parameter.

| Minimum distance, x (oxDNA units, 0.85 nm) | *m* |
|---|---|
| 0<x<1 | 0 |
| 1<x<2 | 1 |
| 2<x<4 | 2 |
| 4<x<8 | 3 |
| 8<x | 4 |

**Table S11:** Windows used in umbrella sampling for identifying free energy differences between the two states in PD-PAINT.

| Window index | Minimum distance range, *m* | Number of bonds, $n_{B.}$ | S1 stem-loop formation | S1S2 binding |
|---|---|---|---|---|
| 0 | 3,4 | 0 | | ✓ |
| 1 | 2,3 | 0 | ✓ | ✓ |
| 2 | 1,2 | 0 | ✓ | ✓ |
| 3 | 0,1 | 0 | ✓ | ✓ |
| 4 | 0,1 | 0,1 | ✓ | ✓ |
| 5 | 0 | 1,4 | ✓ | ✓ |
| 6 | 0 | 3,6 | ✓ | ✓ |
| 7 | 0 | 5,8 | ✓ | ✓ |
| 8 | 0 | 7,10 | ✓ | ✓ |
| 9 | 0 | 9,12 | ✓ | ✓ |
| 10 | 0 | 11,14 | ✓ | ✓ |
| 11 | 0 | 13,16 | | ✓ |
| 12 | 0 | 15,18 | | ✓ |
| 13 | 0 | 17,19 | | ✓ |
| 14 | 0 | 19,20 | | ✓ |

**Table S12:** Umbrella sampling windows for S1-imager hybridization expressed in terms of the 2 reaction coordinates. * Indicates that this window was only used for the case of the closed S1 loop, with a bias to accentuate probability of observing binding events.

| Window | $b$ | $d$ |
|---|---|---|
| B | $0 \leq b \leq 2$ | $0 \leq d \leq 1$ |
| F | $1 \leq b \leq 9$ | 0 |
| S | 0 | $0 \leq d \leq 4$ |
| I | 0 | $4 \leq d \leq 7$ |
| L* | $1 \leq b \leq 9$ | 0 |

**Supplementary Figures**

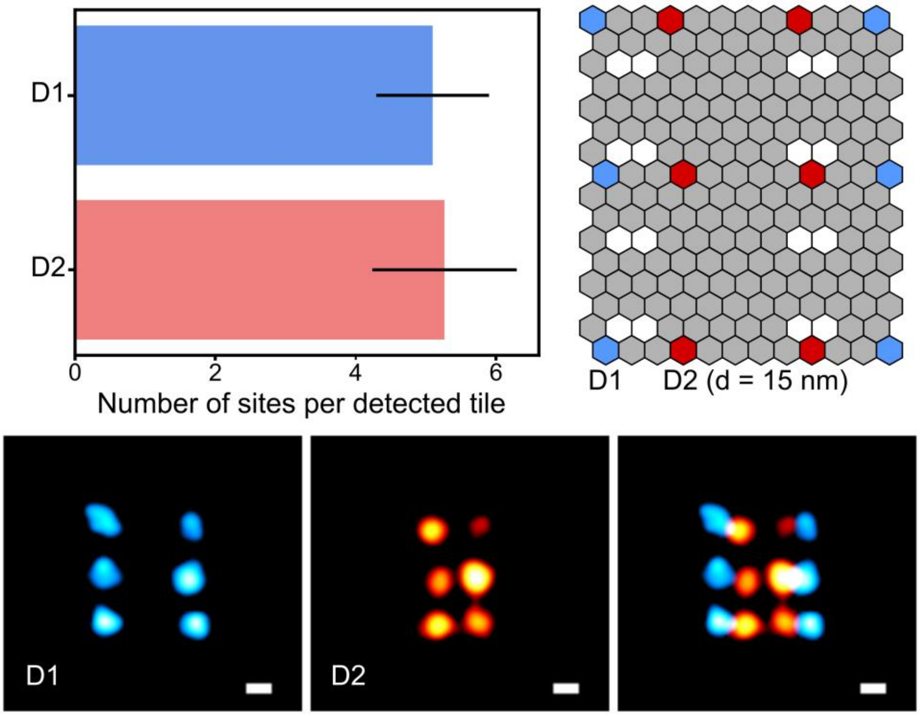

**Figure S1:** Validation of D1 and D2 overhangs as accessible attachment sites on DNA origami. Top: the average number of sites counted per detected origami tile in individual experiments is 5.1 ±0.8 and 5.3 ± 1.0 sites for D1 and D2 markers, respectively (mean ± SD, n = 490 and 400 origami tiles for D1 and D2), equivalent to an addressability of 85% and 88%. Note that while D1 sites were directly targeted with P1 imagers, D2 sites were not probed directly, but first labeled with strand S3, which was then imaged using its P1 docking site. Bottom: example Gaussian rendered images of D1 sites (left) D2 sites (right) and an overlay of these example tiles (right), showing that the detected sites correctly mirror the expected pattern from the design of the origami tile. Scale bars: 20 nm.

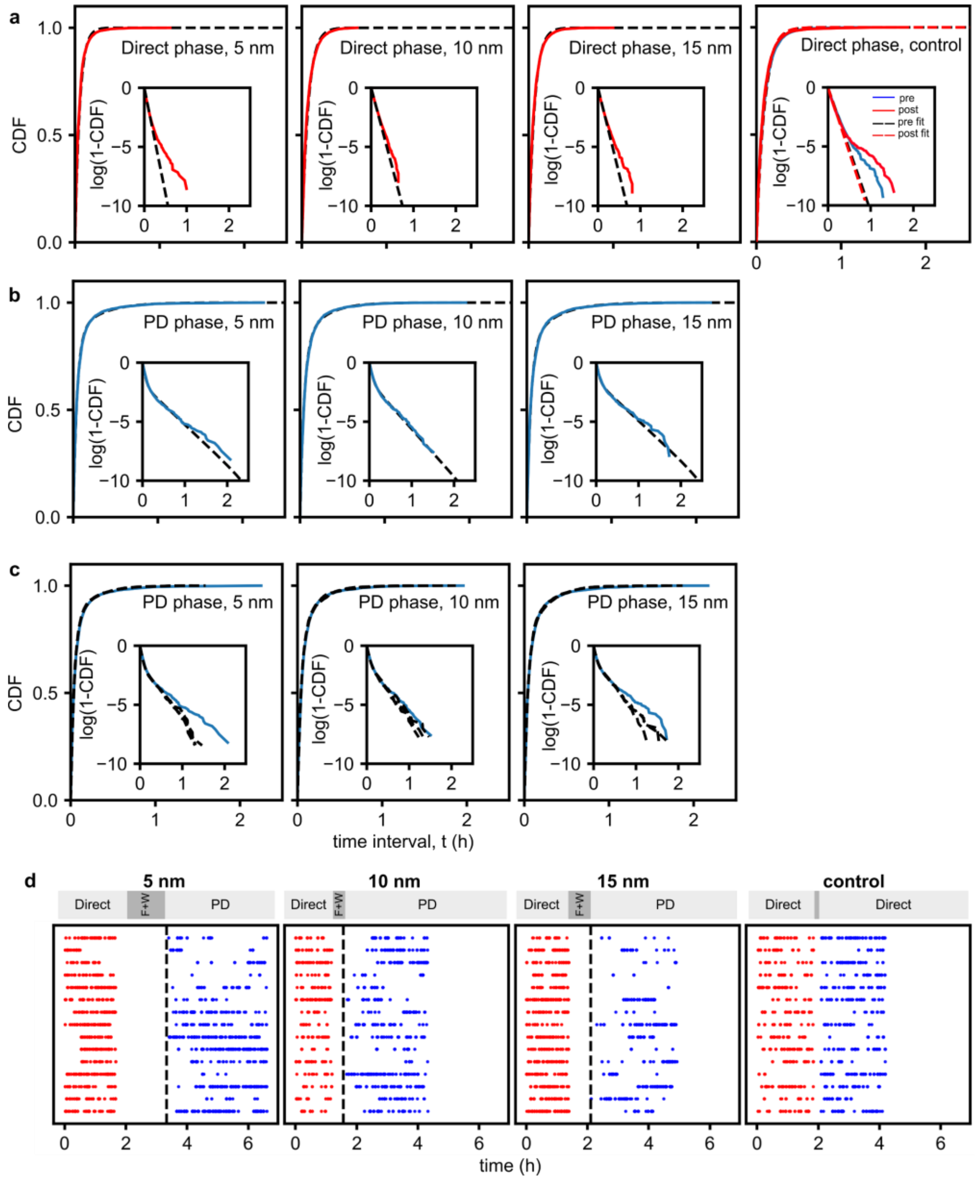

**Figure S2:** Analytical and numerical dark-time cumulative distribution functions (CDF) experimentally derived from the blinking kinetics of single sites on origami test tiles (Fig. 3 and 4). **a:** Dark-time CDFs recorded in the direct phases of PD-PAINT experiments for origami tiles with $d$ = 5 nm, 10 nm and 15 nm, and for control experiments in both the pre- and post-wash direct phases (Fig. 3). Experimental data are shown as solid lines and exponential fits (Eq S4) as dashed lines. The fitted kinetic rates of imager binding to the docking motif are shown in Table S4. **b:** Dark-time CDFs recorded in the PD phases of PD-PAINT experiments at all tested values of $d$ (solid lines) and hyper-exponential analytical fit (dashed lines, Eqs S5-S8). The extracted kinetic parameters for S1-S2 binding/unbinding ($k_d, k_u$) and imager/docking binding ($k_{on}c$) are shown in Table S3. **c:** Data in panel **b** fitted to the outcome of explicit Markov models. Dashed lines indicate, for each plot, 3 numerically generated CDFs with optimized kinetic parameter. Optimized $k_u$ and $k_d$ parameters are shown in Table S5. Insets in all panels show empirical CDFs and theoretical CDFS in log scale. Note that in all CDFs distributions there are slightly more dark events than would be predicted by an exponential model, although this is substantially more pronounced for the PD phases of PD paint experiments, where deviation from the exponential behavior occurs at much shorter timescale. **d:** Example event time traces for the four experiments analyzed in panels **a**-**c**. Symbols indicate individual blinks, while different rows correspond to different sites on the origami. Events for the direct and PD phases are shown in red and blue, respectively. The two experimental phases are separated by functionalization and washing steps (F+W), as discussed in the SI methods and main text (Fig. 3a). Events that might occur during washing/functionalization are excluded from the statistical analysis and not shown in the plots. The dashed line indicates the addition of shield removal strand R. Note that in the control experiment functionalization is not performed, nor is R added (Fig. 3b). The experiment is terminated after between 4 and 6.5 hours, indicated by the sudden end in collection of bright events.

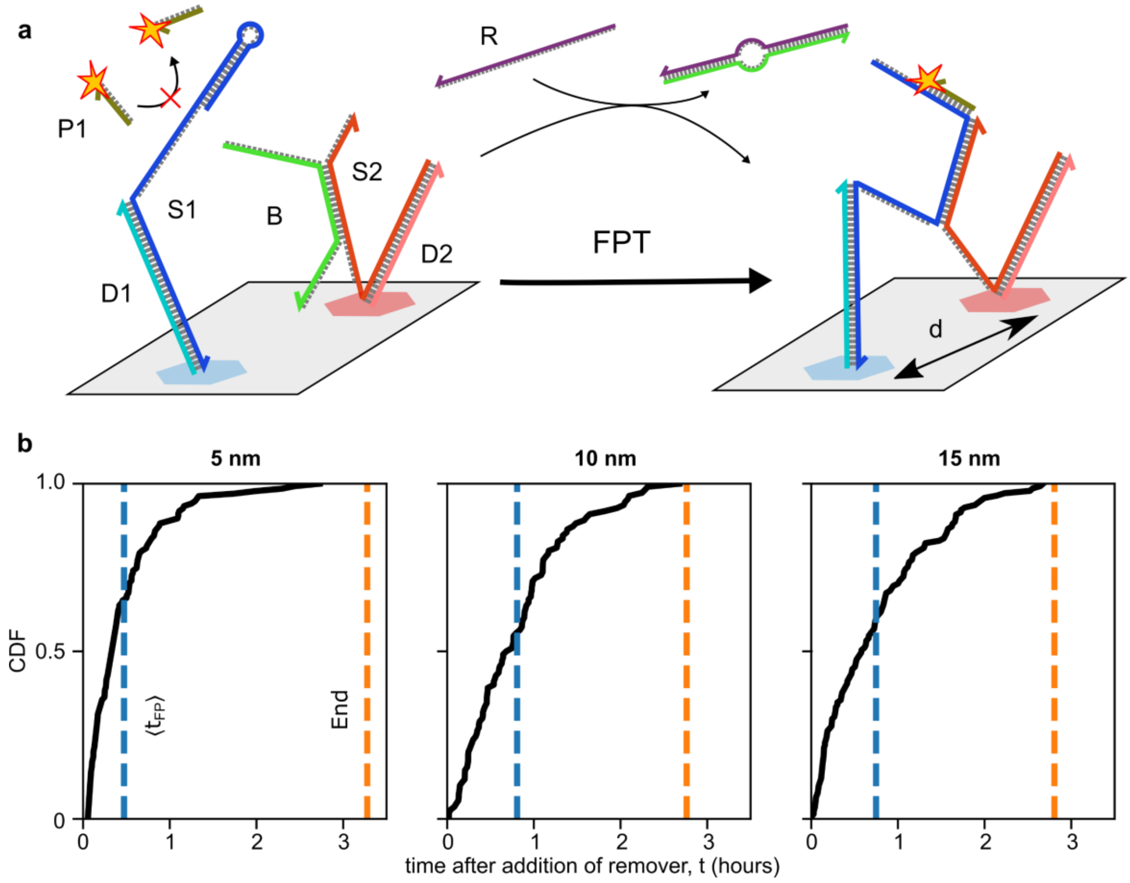

**Figure S3:** Statistic analysis of event first passage time for PD-PAINT as determined on origami tiles. **a:** Schematic indicating the time interval we define as the first passage time $t_{fp}$, namely from the moment the remover strand R is added to the detection of the first bright event on each given origami site (see also Fig. 1). **b:** Cumulative distribution function of $t_{fp}$ as experimentally determined for origami tiles with D1-D2 spacing $d$ = 5, 10 and 15 nm. Blue dashed lines indicate the mean first passage time $<t_{fp}>$ ; orange dashed lines indicate the end of the experiment. Note that in all cases $<t_{fp}>$ is significantly shorter than the duration of the experiment, indicating that the vast majority of the viable sites on the origami should produce blinks over experimental timescales. Note also that the timescale of the toeholding reaction leading to the displacement of the shield strand B after the addition of R is negligible compared to the measured $<t_{fp}>$. The rate of strand displacement can be approximated as $k_{on}[R]$, where $k_{on}$ is the oligo-oligo hybridization rate constant and [R] the shield removal concentration.[23] Using the literature value of $k_{on} \approx 10^6\ M^{-1}s^{-1}$ and $[R]\ >\ 500$ nM we find a strand-displacement rate > 0.5 $s^{-1}$.

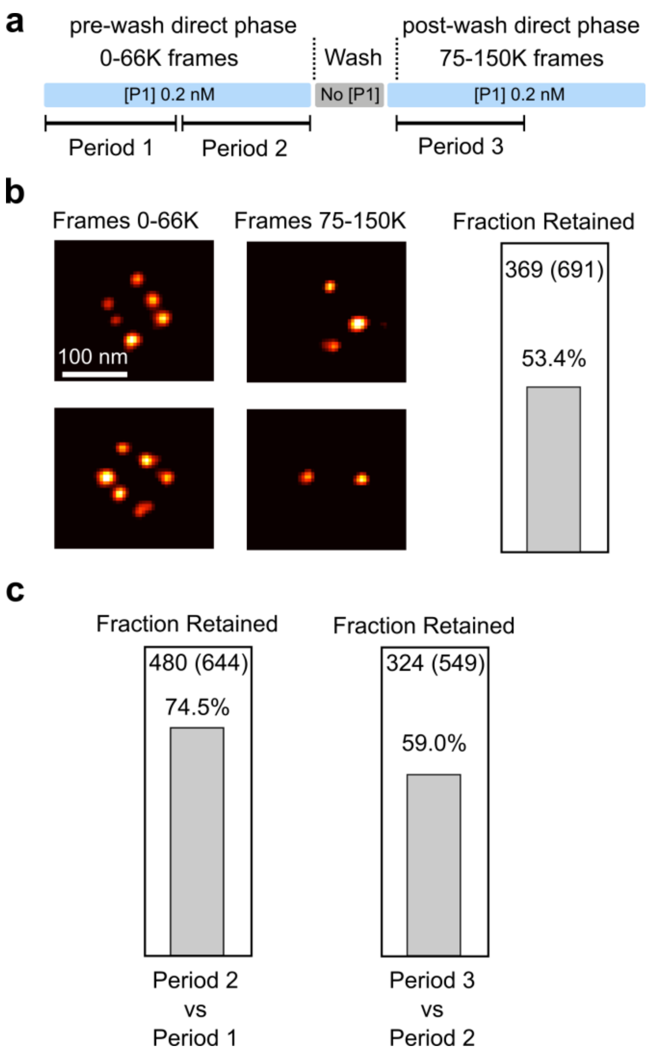

**Figure S4:** Evaluation of loss during conventional DNA-PAINT experiments on origami test tiles. **a**: Protocol of a control experiment where P1 imagers are directly binding to a docking motif on attachment sites D1. As shown graphically in Fig. 3. The origami is first imaged for 66K frames in the presence of imager P1 (pre-wash direct phase), then the P1 imager is removed by washing and finally again reintroduced so that from 75K frames onward the event rate was constant again (post-wash direct phase). **b**: DNA-PAINT images as rendered from the first 66K frames and from frames 75-151K frames, respectively. Comparison of the detected spots showed that ~53% of sites were retained in the second sequence which was rendered from frames 75-151K. **c**: To estimate if sites were lost during normal imaging or as a result of washing we also rendered 3 images from frame periods 1, 2 & 3 (as indicated in **a**) which each comprise 33K frames. Site retention between periods 1 and 2 was 75% whereas site retention between periods 2 and 3 was 59%. This indicates that a major proportion of the site loss occurs progressively during normal imaging. There may be an additional effect due to the washing procedure, but this appears smaller than the imaging induced site loss. Imaging induced site-loss has been shown to result from chemical DNA modifications due to dye radicals formed as a result of fluorescence excitation by Blumhardt et al.[24]

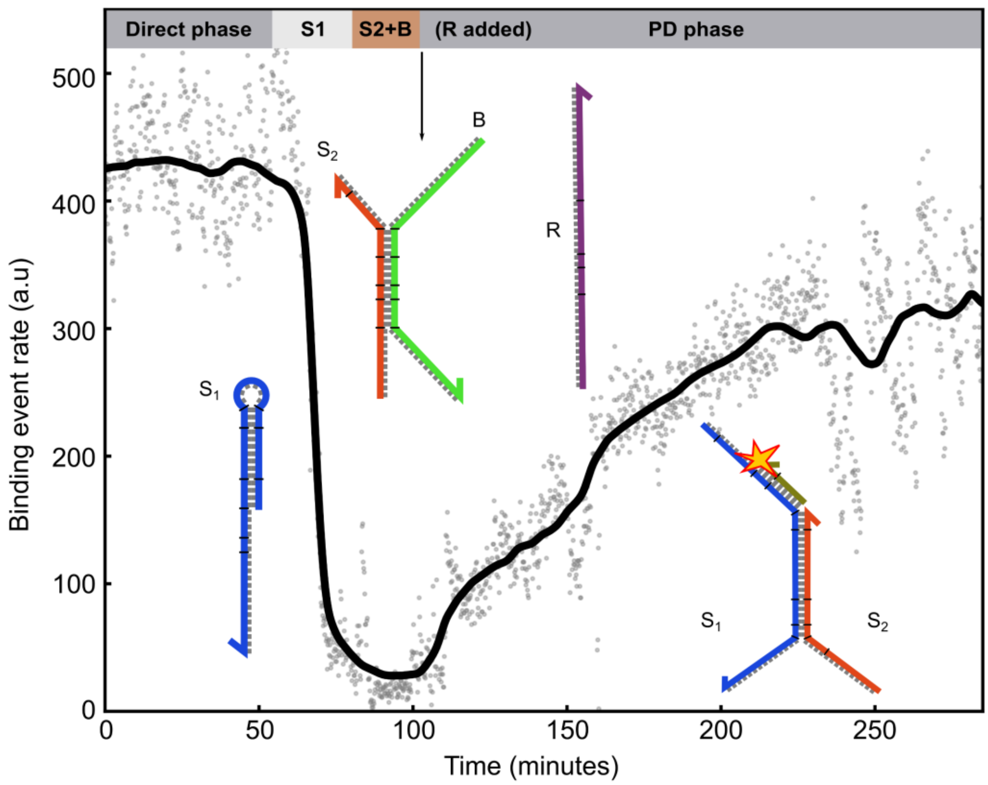

**Figure S5**. Example of an event-rate time trace for an origami PD-PAINT experiment as discussed in the main text (Fig. 1 & 3), here for the case of d = 15 nm. In the initial direct phase D1 was imaged with P1 (2nM). We then added S1 (500 nM) and later S2 (500 nM) pre-mixed with excess B (2 μM) in DNA-PAINT buffer (see experimental methods), allowing for both to bind the respective origami sites (D1 and D2). The addition of the remover strand (500 nM), at arrow location, triggered S1-S2 dimerization and the detection of a PD-PAINT signal (PD phase). Note the progressive increase in event rate which we study with single site resolution (Fig. 4 and Fig. S2). P1 imager concentration was maintained throughout. Individual points indicate raw experimental event data and the black solid line is with a median filter applied.

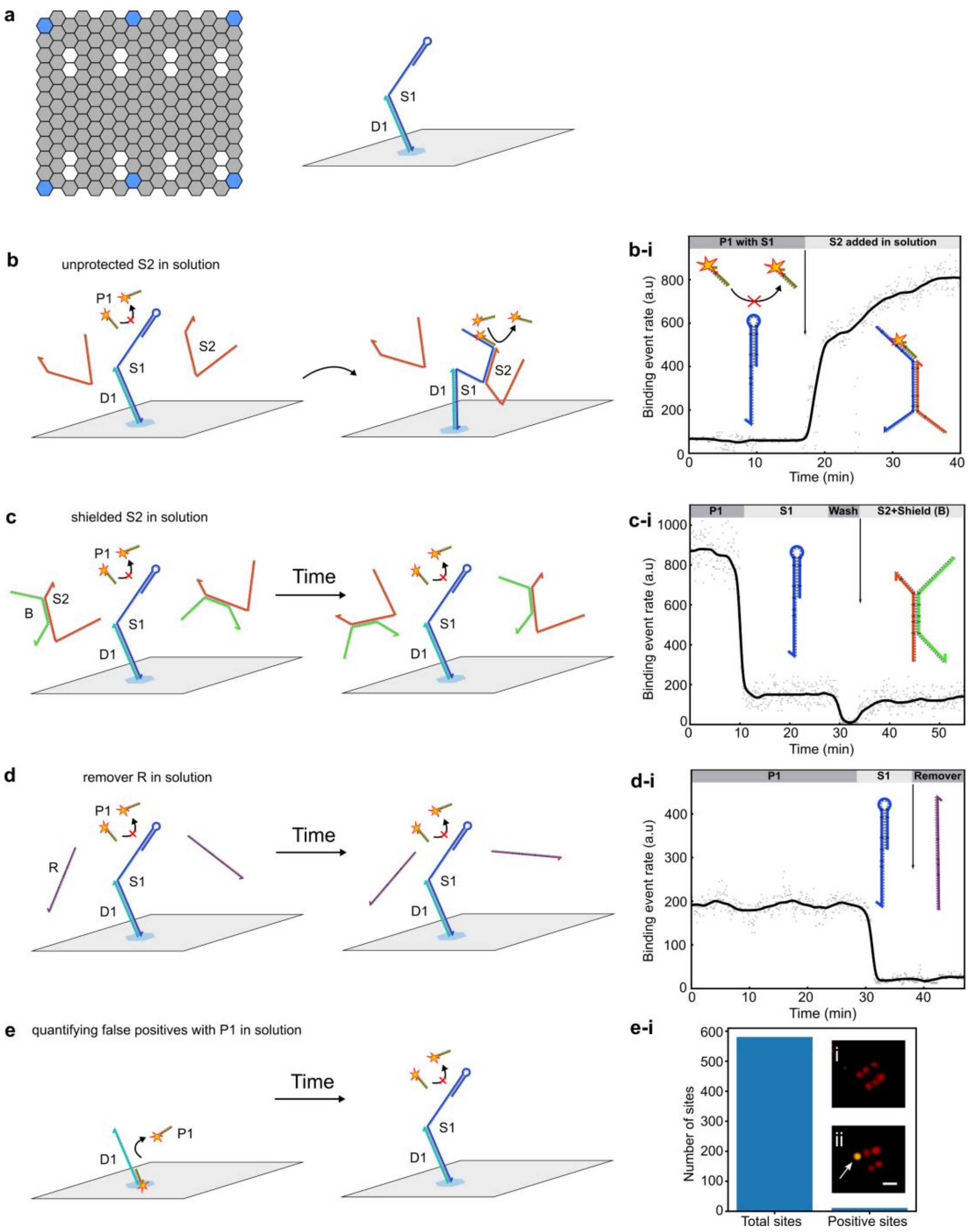

**Figure S6:** Control experiments conducted using synthetic DNA origami samples. **a:** All controls shown here used DNA-origami that only had D1 attachment sites, to which S1 strands could be attached. **b:** Origami initially featuring S1 motifs connected to D1 produced no DNA-PAINT signal from imager P1, given that the docking site on D1 is blocked by S1 and that the docking site on S1 obstructed by loop formation. The addition in solution of large concentrations (~500 nM) of unbound S2 strands, lacking the shield strand B, lead to the formation of S1-S2 dimers, the opening of the S1 loops and the detection of a DNA-PAINT signals, as seen in the time trace of event rates (**b-i**). **c:** If the same experiment as in panel **b** is performed by adding protected S2-B complexes (>500 nM), S1-S2 dimerization is hindered and no DNA-PAINT signal is detected (**c-i**). **d:** Analogous experiment where the shield-remover R is added (>500 nM), also causing no increase in event rates (**d-i**). **e**: Control experiment in which initially D1 is directly targeted with P1, leading to the detection of a DNA-PAINT signal from origami binding sites, followed by the addition of S1 which block the P1 docking domains on D1 for the vast majority of origami sites. In **e-i** we compared the number of sites initially detected by imaging D1 (green spots in the rendered images) with those still visible after introducing S1 (red spots). The percentage of the latter is 1.89%, demonstrating a very low prominence of false positive signals in PD-PAINT. The P1 imager concentration remained constant throughout the control experiments. In event-traces (**b-i**, **c-i**, and **d-i**) individual points indicate raw experimental event data and the black solid line is with a median filter applied. Scale bar: 60 nm.

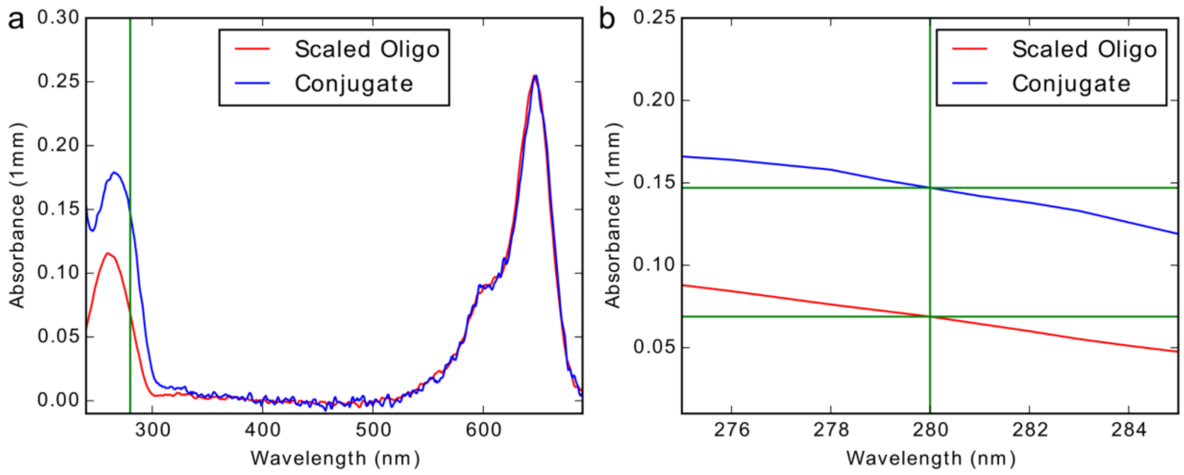

**Figure S7:** Quantification of conjugation efficiency between oligonucleotides and antibodies. **a:** Absorbance measurements of the dye-modified docking oligonucleotide strands were taken prior to conjugation. These were then scaled to match the absorbance peak of the dye used, here Cy5, of the final antibody-oligonucleotide conjugation. **b:** The difference between the absorbance values at 280 nm were attributed to the contribution from the antibody, assuming that all oligonucleotides have a modified dye attached. From this the ratio of oligo-antibody could be calculated, typically obtaining >1-3 dye-oligonucleotides per antibody, see details in SI Methods.

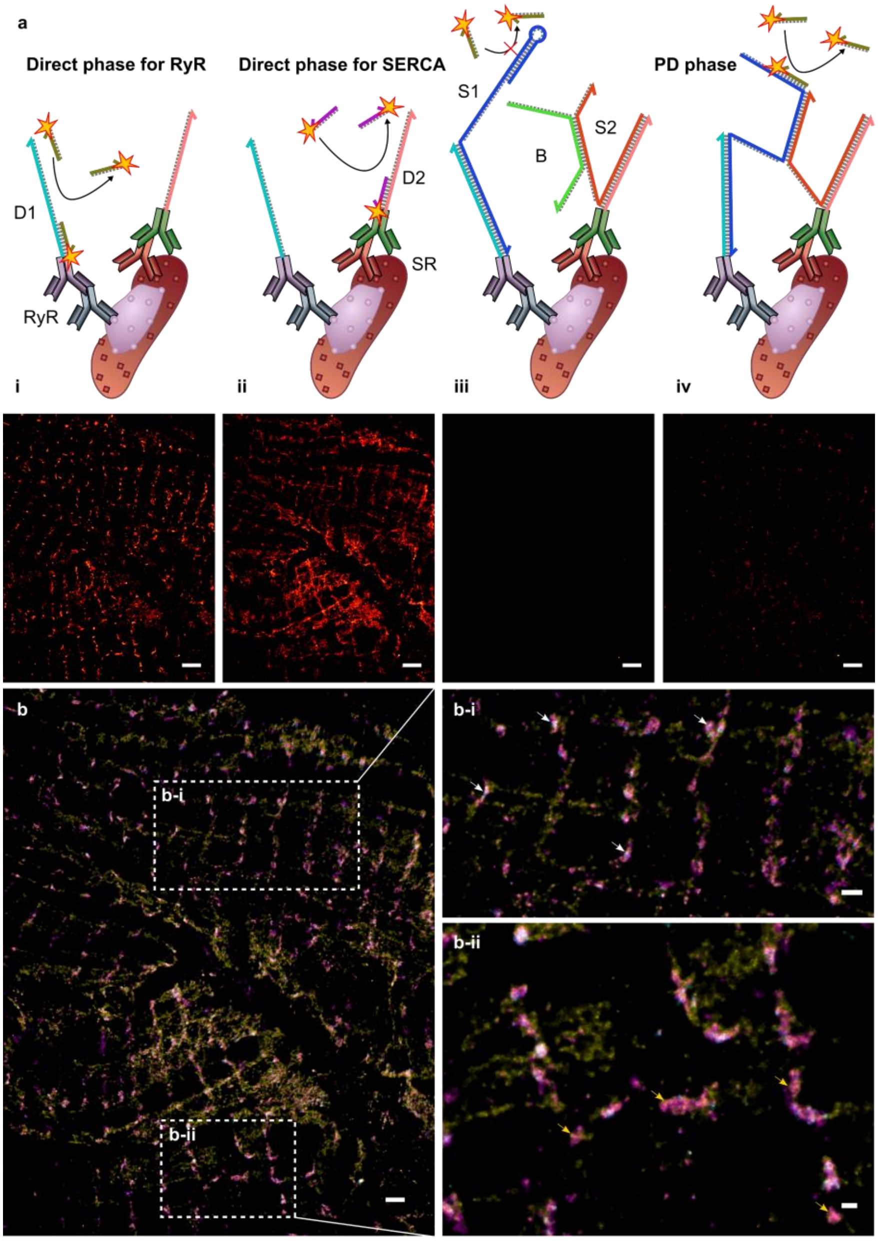

**Figure S8:** PD-PAINT in cardiac tissue sections (not to scale). **a:** Primary and secondary antibody labeling of ryanodine receptors (RyR) using D1 which has a P1 docking site **(i)** and the sarcoplasmic reticulum (SR) protein SERCA with D2 containing a P5 docking site **(ii)**. An Exchange-PAINT protocol allows for the sequential imaging of both targets **(i-ii)**. S1 strands bind to the extended docking sequence D1, preventing P1 imager from interacting with the docking domain, a shielded S2 strand does the same for P5 by binding to D2 site **(iii)**. Removal of the S2 shield allows S1 and S2 probes anchored within close proximity dimerize and the S1 loop. The open S1 loops exposes a docking motif which can be sampled with P1 imagers **(iv)**. Bottom: rendered DNA-PAINT images of the respective steps **(i-iv)**. **b:** An overlay of the DNA-PAINT images from **(a)** (RyR P1 – magenta, SERCA P5 – yellow, PD-PAINT – cyan). PD-PAINT signal typically only appears where both P1 and P5 signal show co-localization (boxes). White arrows **(b-i)** highlight regions with PD-PAINT signal, yellow arrows **(b-ii)** point to regions where co-localization would suggest molecules are in close proximity but do not display PD-PAINT signal, indicating a greater distance between target epitopes at these locations. Frame numbers used to generate super-resolution images: D1 & D2 ~40k frames & open S1 sites ~50k frames. Scale bars: **a** 2 µm, **b** 1 µm, **b-i** 500 nm, **b-ii** 250 nm.

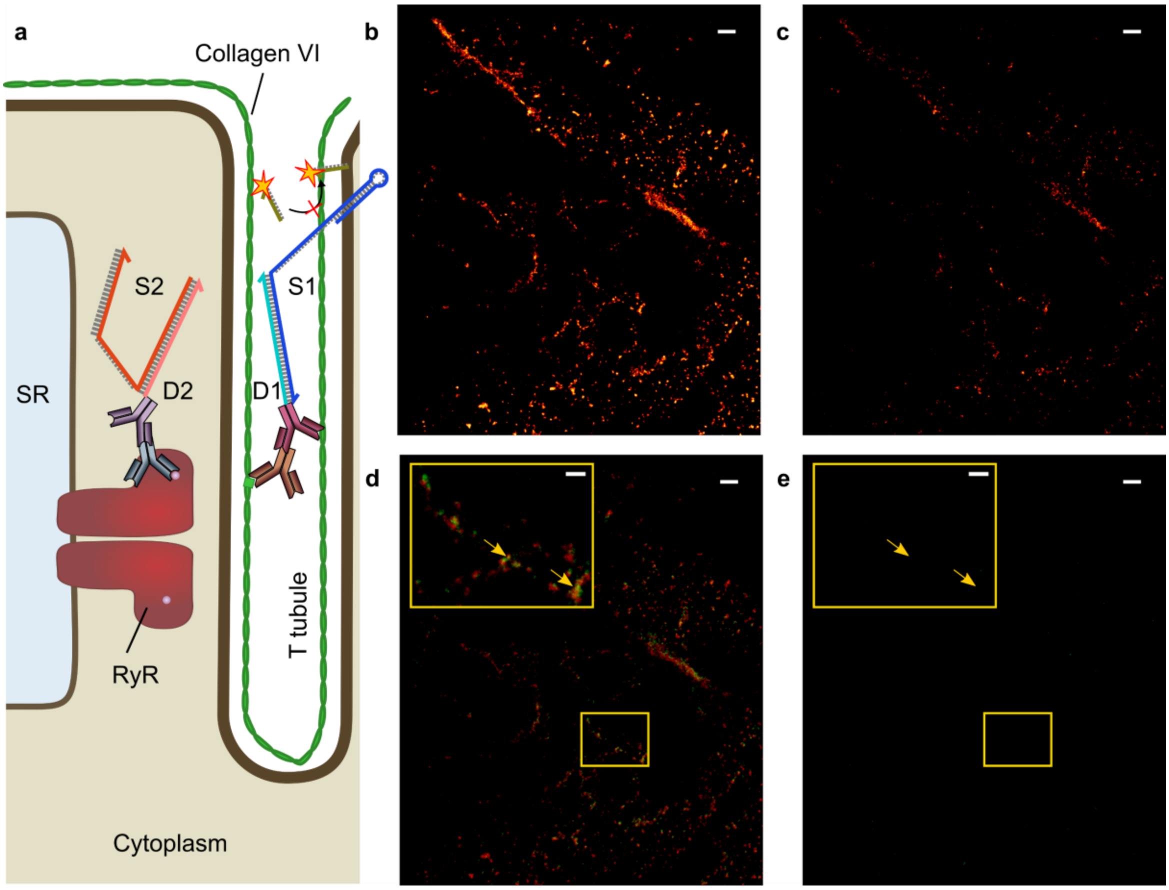

**Figure S9:** Negative control experiment in biological tissue. **a:** Not-to scale schematic of locations of Collagen VI (ColVI)[25] and RyR, including invaginations of the cell membrane referred to as T tubules. The RyR is a membrane protein of the sarcoplasmic reticulum (SR), located inside the cell, while ColVI is located outside. We labeled RyR with D2 and ColVI with D1. **b:** Collagen-VI DNA-PAINT image obtained by targeting D1 with imager P1. **c:** RyR DNA-PAINT image obtained by targeting D2 with P5. **d:** ColVI (red) – RyR (green) overlay. Arrow highlight regions of apparent co-localization. **e**: PD-PAINT data are collected after binding S1 and S2 to D1 and D2 respectively, and removing the shield strand B. The lack of PD-PAINT signal in the rendered image suggests that the distance between ColVI and RyR is too large for S1 and S2 to hybridize and that the apparent co-localization seen in the ColVI and RyR signal overlay **(d)** does not reflect molecular proximity. Frame numbers used to generate super-resolution images: D1, D2 & open S1 sites ~30k frames each. Scale bars: 500 nm, insets 200 nm.

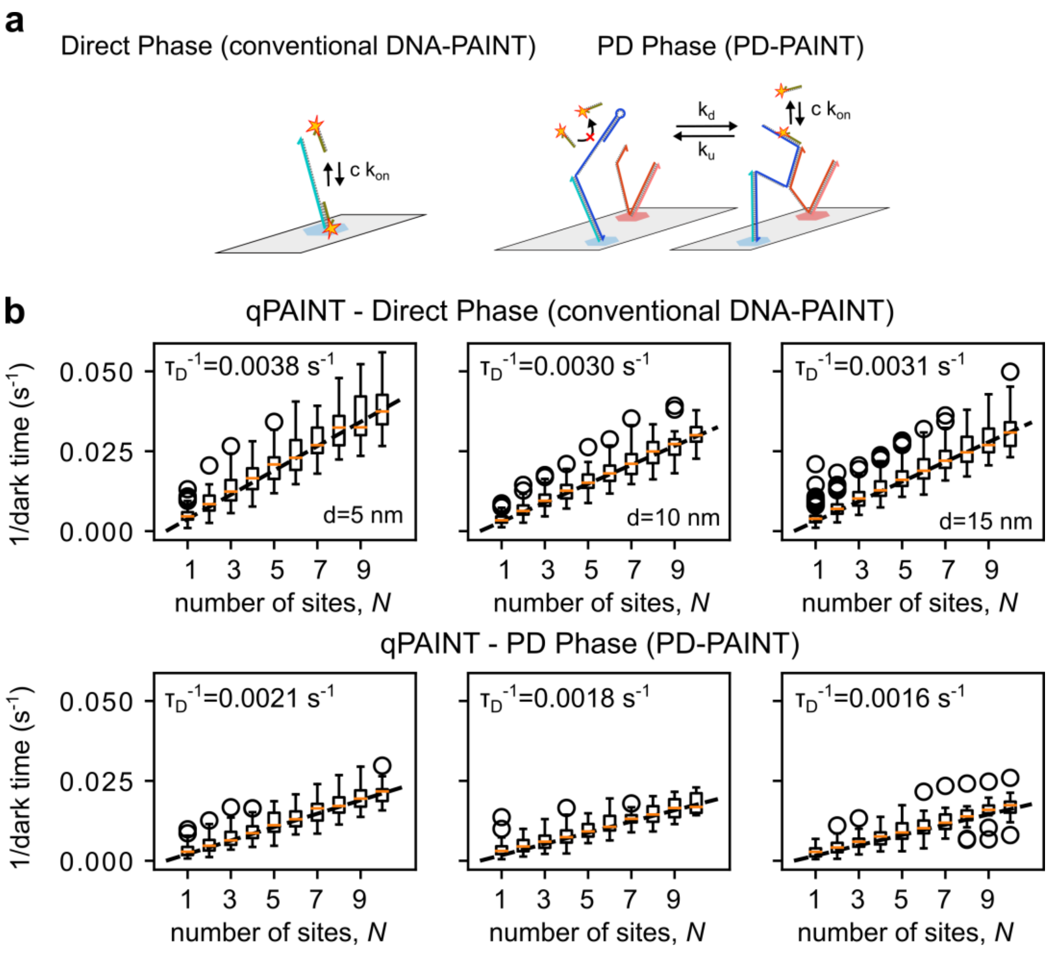

**Figure S10.** qPAINT analysis of dark times between detected imager binding events at single origami docking sites. In qPAINT, the number of docking sites scales linearly with the inverse mean dark time for a given region containing *N* docking sites. The mean dark time can be estimated from the time series of bright events, and allows estimation of the number of docking sites, given a 'calibration' by the inverse mean dark time of a single docking site. **a:** We validated the possibility of performing qPAINT with the PD-PAINT scheme which exhibits long dark times due to transitions between the open and closed S1 loop states. **b:** To generate a time series representative of *N* docking sites, *N* single docking site time series (see Fig. 4) were merged, as discussed in the experimental methods. This procedure was carried out for both the direct phases, in which "conventional" DNA-PAINT is performed by targeting D1 with imager P1, and the PD phase, in which PD-PAINT signal is detected (see **a**). The inverse of the mean dark time as a function of *N* fall approximately on a straight line, both for conventional DNA-PAINT (top row), and for PD-PAINT data from time series on origami (bottom row), for *d* = 5, 10 and 15 nm. However, the gradient in the relationship between *N* and the inverse dark time, equal to the single-site inverse mean d dark time $\tau_D^{-1}$, is lower for PD-PAINT than for conventional DNA-PAINT (taken at the same imager concentration). This is a consequence of long dark events (see scheme in **a**) which correspond to the S1 loop transiently closing. qPAINT analysis of PD-PAINT data can be thus be carried out virtually unmodified from conventional DNA-PAINT qPAINT analysis, with the caveat that calibration must be carried out on PD-PAINT data. The variation in calibration factors $\tau_D^{-1}$ for different distances *d* is small and can be largely explained by small differences in imager concentration.

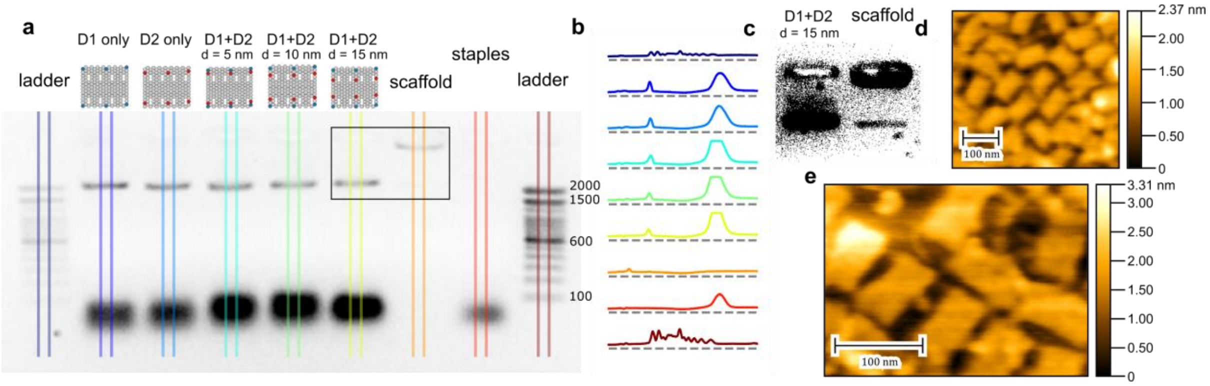

**Figure S11: a:** Agarose gel (1.5%) of annealed DNA origami, free staples and scaffold. Annealed DNA origami lanes consist of the origami as the slow migrating major product, with excess staples also visible. The first and the last lane contain 100 bp ladders (ThermoScientific), where several reference bands have been labeled. **b:** Intensities of cross sections through the gel image illustrating origami peaks at the same locations for the five origami. **c:** A thresholded image of the region circumscribed by the black rectangle, demonstrated the non-aggregated scaffold migrating slightly faster than the DNA origami, similarly observed by Jungmann and coworkers for their similar RROs.[26] **d:** AFM micrograph of one of the tested origami designs, namely the one bearing only 6xD1 overhands, confirming a size of approximately 70 nm x 90 nm, and a thickness of 2 nm, comparable with the width of a double helix. **e:** AFM micrographs of a RRO tile lacking overhangs. Excess scaffold is routed through one edge and can be clearly visualized here as a central nodule emanating from one side.

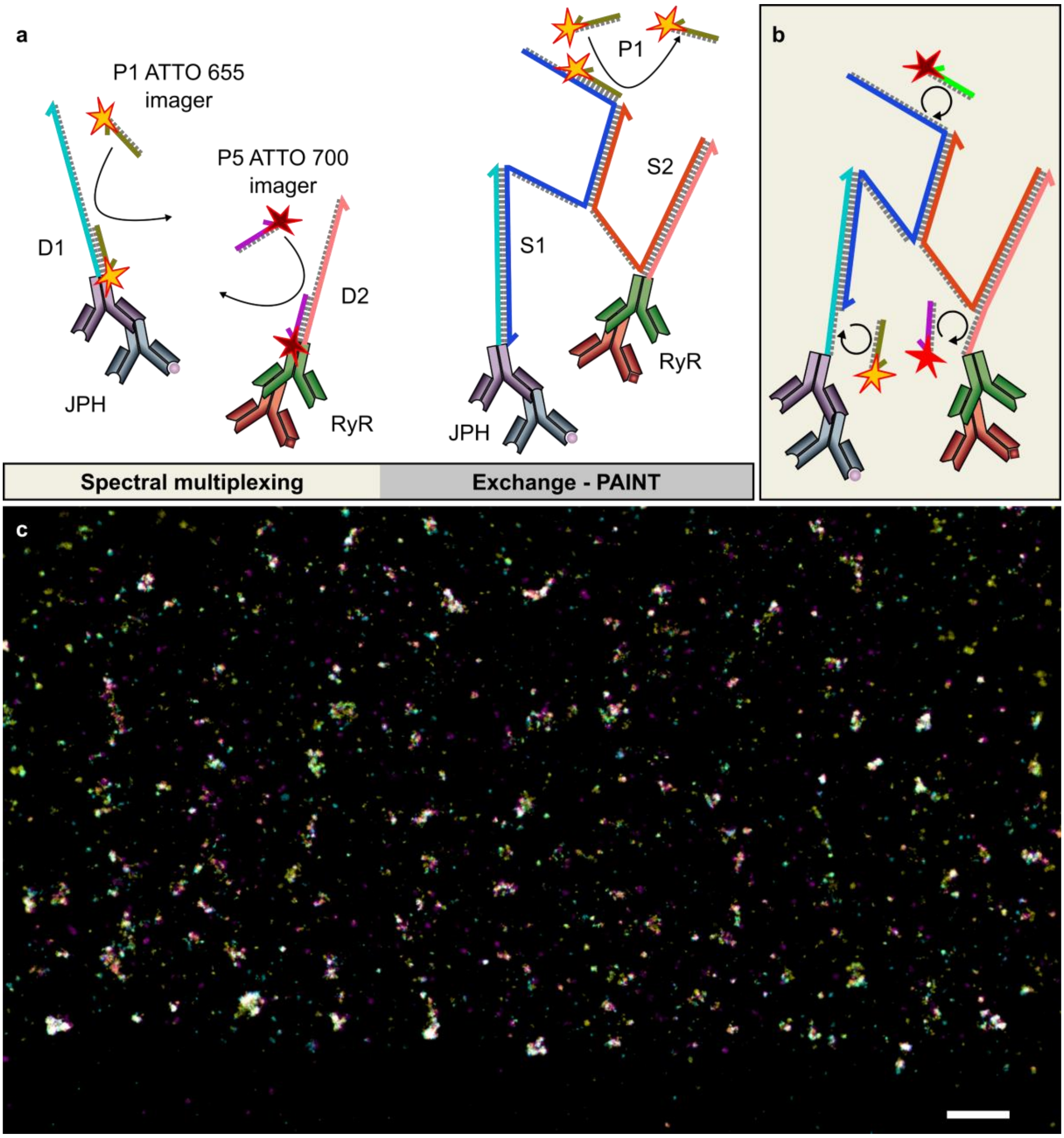

**Figure S12:** Multichannel PD-PAINT imaging of two target proteins. **a:** Not-to-scale sketches of RyR and JPH immuno-labeled with strands D2 and D1, respectively. To perform simultaneous DNA-PAINT imaging of both targets by spectral multiplexing P1 and P5 imagers, targeting D1 and D2, have been modified to carry ATTO 655 and ATTO 700 dyes, respectively. Both imagers are simultaneously excited by a single laser source and the detection path is spectrally split. For details on the optical setup see Baddeley et al[8]. After the spectral multiplexing measurement S1 and S2 strands can be attached to D1 and D2 as discussed in the main text, and the S2 is unprotected to allow S1-S2 dimerization and the detection of a PD-PAINT signal by sampling with P1. **b:** Proposed use of multiplexed PD-PAINT for imaging exposed P1 and P5 docking sites on (modified) D1 and D2 strands at the same time as any open S1 domains via spectral or temporal multiplexing. Such a design has the benefit that all labeling and attachment reactions can be completed before beginning imaging experiments. **c:** A Ratio-PD-PAINT biological experiment as depicted in **(a)**. RyR (magenta) and JPH (yellow) data acquired simultaneously and subsequent open loop domains (cyan) via conventional Exchange–PAINT. Scale bar: 1µm.

**References**

(1) Schnitzbauer, J.; Strauss, M. T.; Schlichthaerle, T.; Schueder, F.; Jungmann, R. Super-Resolution Microscopy with DNA-PAINT. *Nat. Protoc.* **2017**, *12* (6), 1198–1228. https://doi.org/10.1038/nprot.2017.024.

(2) Nečas, D.; Klapetek, P. Gwyddion: An Open-Source Software for SPM Data Analysis. *Cent. Eur. J. Phys.* **2012**, *10* (1), 181–188. https://doi.org/10.2478/s11534-011-0096-2.

(3) Jungmann, R.; Steinhauer, C.; Scheible, M.; Kuzyk, A.; Tinnefeld, P.; Simmel, F. C. Single-Molecule Kinetics and Super-Resolution Microscopy by Fluorescence Imaging of Transient Binding on DNA Origami. *Nano Lett.* **2010**, *10* (11), 4756–4761. https://doi.org/10.1021/nl103427w.

(4) Jayasinghe, I.; Clowsley, A. H.; Lin, R.; Baddeley, D.; Michele, L. Di; Soeller, C.; Jayasinghe, I.; Clowsley, A. H.; Lin, R.; Lutz, T.; Harrison, C.; Green, E.; Baddeley, D.; Michele, L. Di; Soeller, C. True Molecular Scale Visualization of Variable Clustering Properties of Ryanodine Receptors. *Cell Rep.* **2018**, *22* (2), 557–567. https://doi.org/10.1016/j.celrep.2017.12.045.

(5) Jayasinghe, I. D.; Cannell, M. B.; Soeller, C. Organization of Ryanodine Receptors, Transverse Tubules, and Sodium-Calcium Exchanger in Rat Myocytes. *Biophys. J.* **2009**, *97* (10), 2664–2673. https://doi.org/10.1016/j.bpj.2009.08.036.

(6) Crossman, D. J.; Hou, Y.; Jayasinghe, I.; Baddeley, D.; Soeller, C. Combining Confocal and Single Molecule Localisation Microscopy: A Correlative Approach to Multi-Scale Tissue Imaging. *Methods* **2015**, *88*, 98–108. https://doi.org/10.1016/j.ymeth.2015.03.011.

(7) Oort, R. J. Van; Garbino, A.; Wang, W.; Dixit, S. S.; Landstrom, A. P.; Gaur, N.; Almeida, A. C. De; Skapura, D. G.; Rudy, Y.; Burns, A. R.; Ackerman, M. J.; Wehrens, X. H. T. Disrupted Junctional Membrane Complexes and Hyperactive Ryanodine Receptors After Acute Junctophilin Knockdown in Mice. *Circulation* **2011**, *123* (9), 979–988. https://doi.org/10.1161/CIRCULATIONAHA.110.006437.

(8) Baddeley, D.; Crossman, D.; Rossberger, S.; Cheyne, J. E.; Montgomery, J. M.; Jayasinghe, I. D.; Cremer, C.; Cannell, M. B.; Soeller, C. 4D Super-Resolution Microscopy with Conventional Fluorophores and Single Wavelength Excitation in Optically Thick Cells and Tissues. *PLoS One* **2011**, *6* (5), 1–10. https://doi.org/10.1371/journal.pone.0020645.

(9) Lutz, T.; Clowsley, A. H.; Lin, R.; Pagliara, S.; Michele, L. Di. Versatile Multiplexed Super-Resolution Imaging of Nanostructures by Quencher-Exchange-PAINT. *Nano Res.* **2018**, 1–14. https://doi.org/https://doi.org/10.1007/s1227.

(10) Mcgorty, R.; Kamiyama, D.; Huang, B. Active Microscope Stabilization in Three Dimensions Using Image Correlation. *Opt. Nanoscopy* **2013**, *2* (3), 1–7. https://doi.org/https://doi.org/10.1186/2192-2853-2-3.

(11) Baddeley, D.; Cannell, M. B.; Soeller, C. Visualization of Localization Microscopy Data. *Microsc. Microanal.* **2010**, *16* ((1)), 64–72. https://doi.org/https://doi.org/10.1017/S143192760999122X.

(12) Jungmann, R.; Avendaño, M. S.; Dai, M.; Woehrstein, J. B.; Agasti, S. S.; Feiger, Z.; Rodal, A.; Yin, P. Quantitative Super-Resolution Imaging with QPAINT. *Nat. Methods* **2016**, *13* (5), 439–442. https://doi.org/10.1038/nmeth.3804.

(13) Böger, C.; Hafner, A.-S.; Schlichthärle, T.; Strauss, M. T.; Malkusch, S.; Endesfelder, U.; Jungmann, R.; Schuman, E. M.; Heilemann, M. Super-Resolution Imaging and Estimation of Protein Copy Numbers at Single Synapses with DNA-Point Accumulation for Imaging in Nanoscale Topography. *Neurophotonics* **2019**, *6* (03), 1. https://doi.org/10.1117/1.nph.6.3.035008.

(14) Beyer, J.; Nielsen, B. Predator Foraging in Patchy Environments: The Interrupted Poisson Process (IPP) Model Unit. *Dana* **1996**, *11* (2), 65–130.

(15) Ouldridge, T. E.; Louis, A. A.; Doye, J. P. K. DNA Nanotweezers Studied with a Coarse-Grained Model of DNA. *Phys. Rev. Lett.* **2010**, *104* (17), 1–4. https://doi.org/10.1103/PhysRevLett.104.178101.

(16) Ouldridge, T. E.; Louis, A. A.; Doye, J. P. K. Structural, Mechanical, and Thermodynamic Properties of a Coarse-Grained DNA Model. *J. Chem. Phys.* **2011**, *134* (8). https://doi.org/10.1063/1.3552946.

(17) Srinivas, N.; Ouldridge, T. E.; Šulc, P.; Schaeffer, J. M.; Yurke, B.; Louis, A. A.; Doye, J. P. K.; Winfree, E. On the Biophysics and Kinetics of Toehold-Mediated DNA Strand Displacement. *Nucleic Acids Res.* **2013**, *41* (22), 10641–10658. https://doi.org/10.1093/nar/gkt801.

(18) Shi, Z.; Castro, C. E.; Arya, G. Conformational Dynamics of Mechanically Compliant DNA Nanostructures from Coarse-Grained Molecular Dynamics Simulations. *ACS Nano* **2017**, *11* (5), 4617–4630. https://doi.org/10.1021/acsnano.7b00242.

(19) Torrie, G. M Valleau, J. . Nonphysical Sampling Distributions in Monte Carlo Free-Energy Estimation: Umbrella Sampling. *J. Comput. Phys.* **1977**, *23* (2), 187–199.

(20) Whitelam, S.; Geissler, P. L. Avoiding Unphysical Kinetic Traps in Monte Carlo Simulations of Strongly Attractive Particles. *J. Chem. Phys.* **2007**, *127* (15). https://doi.org/10.1063/1.2790421.

(21) Kumar, S.; Rosenberg, J. M.; Bouzida, D.; Swendsen, R. H.; Kollman, P. A. THE Weighted Histogram Analysis Method for Free-Energy Calculations on Biomolecules. I. The Method. *J. Comput. Chem.*

**1992**, *13* (8), 1011–1021. https://doi.org/10.1002/jcc.540130812.
(22) Zhang, J. X.; Fang, J. Z.; Duan, W.; Wu, L. R.; Zhang, A. W.; Yordanov, B.; Petersen, R.; Phillips, A.; Zhang, D. Y. Predicting DNA Hybridization Kinetics from Sequence. *Nat. Chem.* **2018**, *10* (1), 91–98. https://doi.org/10.1038/nchem.2877.
(23) Zhang, D. Y.; Winfree, E. Control of DNA Strand Displacement Kinetics Using Toehold Exchange. *J. Am. Chem. Soc.* **2009**, *131* (47), 17303–17314. https://doi.org/10.1021/ja906987s.
(24) Blumhardt, P.; Stein, J.; Mücksch, J.; Stehr, F.; Bauer, J.; Jungmann, R.; Schwille, P. Photo-Induced Depletion of Binding Sites in DNA-Paint Microscopy. *Molecules* **2018**, *23* (12), 1–27. https://doi.org/10.3390/molecules23123165.
(25) Crossman, D. J.; Shen, X.; Ju, M.; Munro, M.; Hou, Y.; Middleditch, M.; Shrestha, D.; Li, A.; Lal, S.; Remedios, C. G.; Baddeley, D.; Ruygrok, P. N.; Soeller, C. Increased Collagen within the Transverse Tubules in Human Heart Failure. *Cardiovasc. Res.* **2017**, *113* (8), 879–891. https://doi.org/10.1093/cvr/cvx055.
(26) Schnitzbauer, J.; Strauss, M. T.; Schlichthaerle, T.; Schueder, F.; Jungmann, R. Super-Resolution Microscopy with DNA-PAINT. *Nat. Protoc.* **2017**, *12* (6), 1198–1228. https://doi.org/10.1038/nprot.2017.024.
